# Gradients of receptor expression in the macaque cortex

**DOI:** 10.1101/2021.02.22.432173

**Authors:** Sean Froudist-Walsh, Ting Xu, Meiqi Niu, Lucija Rapan, Daniel S. Margulies, Karl Zilles, Xiao-Jing Wang, Nicola Palomero-Gallagher

## Abstract

Dynamics and functions of neural circuits depend on synaptic interactions mediated by receptors. Therefore, a comprehensive map of receptor organization is needed to understand how different functions may emerge across distinct cortical regions. Here we use *in-vitro* receptor autoradiography to measure the density of 14 neurotransmitter receptor types in 109 areas of macaque cortex. We integrate the receptor data with other anatomical, genetic and functional connectivity data into a common cortical space. We uncovered a principal gradient of increasing receptor expression per neuron aligned with cortical hierarchy from early sensory cortex to higher cognitive areas. A second gradient, primarily driven by 5-HT_1A_ receptors, peaks in the anterior and subcallosal cingulate, suggesting that the macaque may be a promising animal model for major depressive disorder. The receptor gradients may enable rapid, reliable information processing in sensory cortical areas and slow, flexible integration of information in higher cognitive areas.

## Introduction

Flexibility is a hallmark of biological intelligence. A key challenge in modern neuroscience is to discover the cellular, molecular and systems architecture that enables the brain to adapt flexibly and appropriately to a rapidly changing world. The creation of nearly complete maps of the brain’s connections across species at the macroscopic (Glasser et al., 2016), mesoscopic (Majka et al., 2020; Markov et al., 2014a; Oh et al., 2014) and microscopic levels (Scheffer et al., 2020; White et al., 1986) has been a major achievement. However, connectivity alone is insufficient to explain neural circuit dynamics underlying brain functions, which depend on the type and timescale of synaptic transmission mediated by transmitter receptors. Therefore, connectomic approaches, which are blind to receptor types, may not be sufficient to understand the computational capabilities of the cortex. The systematic mapping of multiple receptor densities across cortex would provide a crucial link between the molecular and systems organization of the cortex, complementing ongoing efforts to map the connectome.

In comparison to rodents, macaques and humans share a very similar regional and laminar receptor profile (Zilles and Palomero-Gallagher, 2017a). Parallel recent advances in *in-vivo* neuroimaging (Milham et al., 2020, 2018) and mesoscale connectome mapping (Markov et al., 2014a, 2014b, 2013) have increased the translational potential of studies on the macaque brain. Integration of gold-standard neuroanatomy with *in-vivo* measures of cortical structure and function has great promise to help translation across species and scales of neuroscience, but is still in its infancy (Donahue et al., 2016; Froudist-Walsh et al., 2018, 2020; Hayashi et al., 2020; Rapan et al., 2020; Scholtens et al., 2014; Wang et al., 2020). The mapping of precise receptor and anatomical data to a cortical space that is accessible to neuroimaging researchers could dramatically accelerate our understanding across scales of how the brain works, from the synapse to distributed cognitive networks.

While cortical microcircuits share a canonical organization, their properties vary gradually across the cortex in the form of macroscopic gradients (Wang, 2020). So far, little work has been done to compare gradients of distinct anatomical properties. The seemingly complex connectivity structure of the cortex can be well described by a small number of connectivity gradients (Margulies et al., 2016), with patterns of connectivity smoothly changing across the cortex from early sensory areas to peaks in the higher areas of association cortex. Similar understanding of the brain’s large-scale receptor organization is beginning to emerge. In the mouse brain, subcortical neuromodulatory centres have been identified as ‘connector hubs’, that are well placed to influence interactions between networks (Coletta et al., 2020). Large-scale patterns of receptor expression have been recently described in the human brain (Goulas et al., 2021; Zilles and Palomero-Gallagher, 2017b), but it is not known how the pattern of receptor expression may relate to distributed cognitive networks, and the functions they produce.

Here, we measured the density of 14 types of neurotransmitter receptors across 109 areas of macaque cortex. We mapped the data for these 14 receptors, as well as data on neuron density, dendritic tree size, dendritic spines, retrograde tract-tracing of cortical connections and *in-vivo* estimates of cortical microstructure and functional connectivity onto a common cortical space. We find that the receptor architecture of macaque cortex can be well described by a set of low-dimensional gradients. The principal receptor gradient defines a putative cortical hierarchy. Cortical areas high on the gradient had a higher density of receptors per neuron, more dendritic spines per pyramidal cell, larger dendrites, and a lower T1w/T2w ratio, indicative of less myelin. Receptor gradients also aligned with *in-vivo* functional connectivity gradients, suggesting a possible role for neuromodulatory receptors in shifting activity along cortical hierarchies and between higher cognitive networks.

## Results

### Distributions of 14 receptor types across 109 regions of macaque cortex

We first analyzed receptor distribution patterns for 14 receptors throughout the macaque brain using *in-vitro* receptor autoradiography, which enables the quantification of endogenous receptors in the cell membrane through the use of radioactive ligands (Palomero-Gallagher and Zilles, 2018a). Our analysis included three glutamatergic (AMPA, kainate, NMDA), three GABAergic (GABA_A_, GABA_A/BZ_,GABA_B_) and eight neuromodulatory (acetylcholine M_1_, M_2_, M_3_; serotonin 5-HT_1A_, 5-HT_2,_ noradrenaline *α*_1,_ *α*_2_, dopamine D_1_) receptors. In the raw data, several receptors reached highest densities in primary visual cortex (GABA_A_, acetylcholine M_2_, serotonin 5-HT_2_) (Zilles and Palomero-Gallagher, 2017a)(Fig. S1). A distinct set of receptors reached highest densities in parts of the anterior cingulate, including all glutamatergic receptors, GABA_B,_ serotonin 5- HT_1A_, noradrenaline *α*_1_, and dopamine D_1_. M_1_, GABA_A/Bz_ and α_2_ receptors are notable for having high densities in both cingulate cortex and V1. This suggests some shared anatomical patterns of expression across receptors.

For each receptor, except for 5-HT_1A_, the cortical area with the highest density usually contained roughly two to five times as many receptors (per mg protein) as the least dense area (range 1.67-4.7, min 5-HT_2_, max M_2_). The exception was the 5-HT_1A_ receptor, which reached a peak density of 1185 fmol/mg protein in area a24’ab of anterior cingulate cortex, over 17 times the density of 5-HT_1A_ receptors in area V1 (Fig S1). The degree to which neurotransmitters affect neural activity depends on the density of receptors on the cell surface of a neuron (Lu et al., 2001). Receptor autoradiography quantifies the receptors in the cell membrane, but is blind to the neuron density. However, the neuron density also varies by a factor of five across macaque cortex (Fig S1). We reasoned that what matters functionally is receptor expression level per neuron, hence mapped the receptor data and previously published neuron density data (Collins et al., 2010) to the Yerkes 19 cortical surface (Donahue et al., 2016), and estimated the receptor density per neuron across the cortex for all 14 receptor types (Fig. 1).

**Figure 1.**
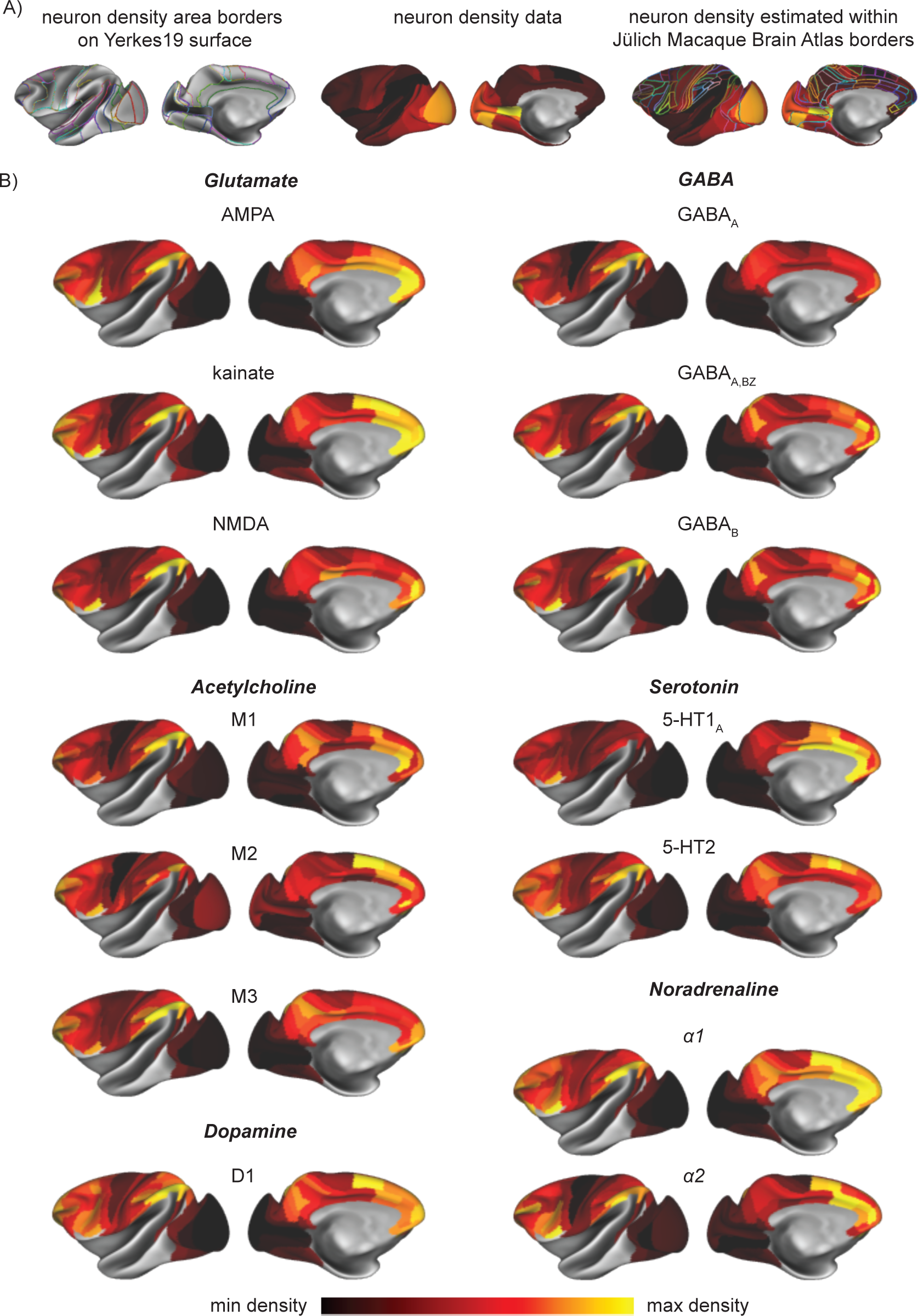
The density of 14 receptors per neuron across macaque cortex. A) Neuron density data from (Collins et al., 2010) was delineated on the cortex and used to normalise receptor data.B) The receptor density per neuron of 14 receptor types assessed with in-vitro receptor autoradiography.

Notably, although the density of several receptors peaks in V1 in the raw data, this is mostly erased when accounting for neuron density. This suggests that the high density of receptors in V1 is largely due to its exceptionally high neuron density (with the possible exception of the M_2_ receptor).

### The principal receptor gradient of macaque cortex

Τo identify the principal patterns of spatial variation across receptors, we performed a principal components analysis (PCA) on the receptor density per neuron data. The first principal component (*principal receptor gradient*), spread from early visual cortex at one end, to association areas of anterior cingulate cortex, orbitofrontal cortex, lateral prefrontal and lateral parietal cortex at the other end. This component alone explained 81% percent of the variance in the receptor data, with the top five principal components sufficient to explain 95% (Fig 2A, Fig. S2, PCs1-5 explain 81.2%, 6.5%, 3.5%, 2.4% and 1.4% respectively). Projection of the data onto the first two principal components (“*receptor space*”) revealed a differentiation of areas into anatomo-functional clusters (Fig 2B), with visual, somatomotor, premotor, parietal, cingulate, prefrontal and orbitofrontal areas occupying distinct sections of the receptor space.

**Figure 2.**
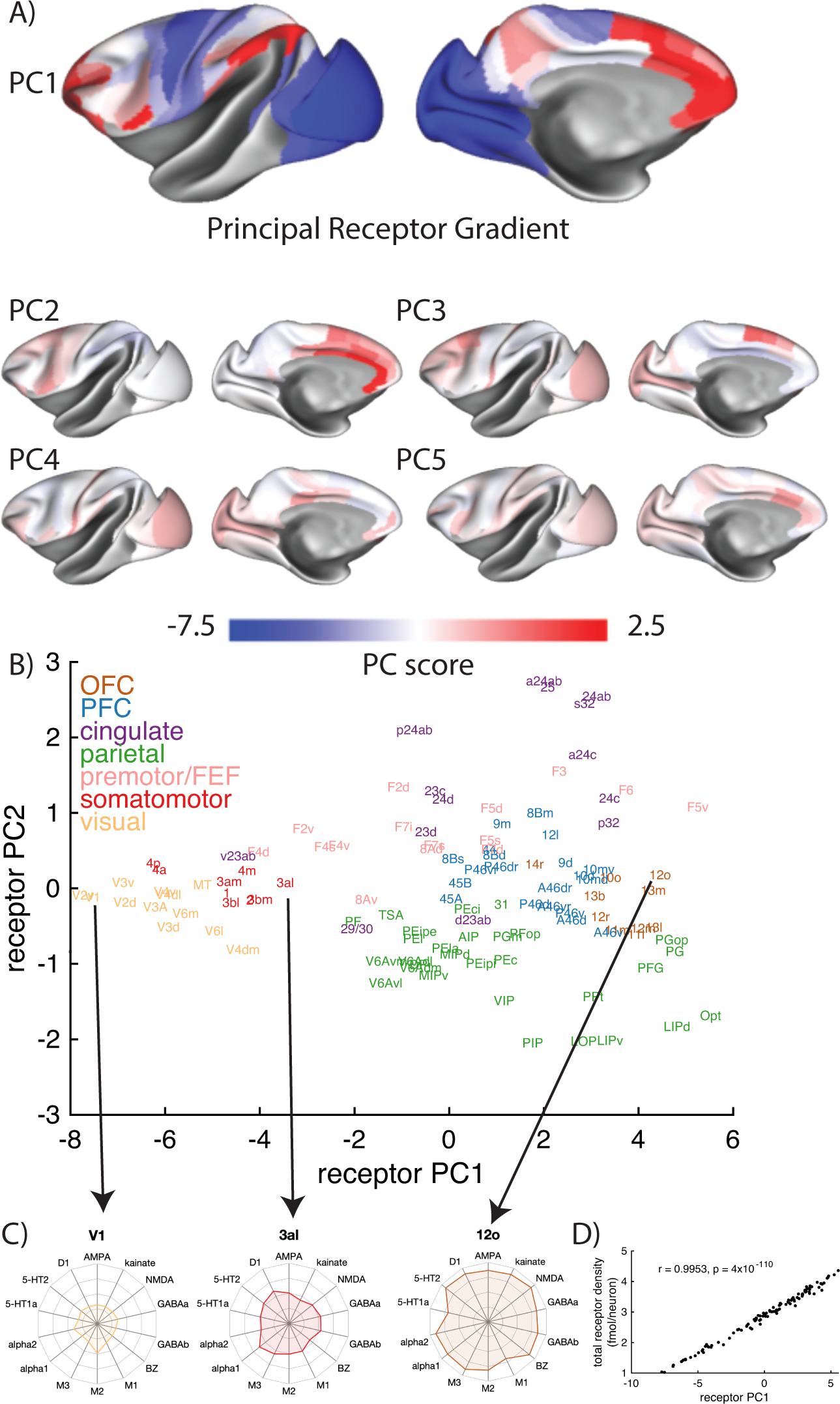
The principal receptor gradient captures total receptor density per neuron across macaque cortex. A) The first five principal components of the receptor per neuron data. B) The projection of brain regions onto the first two principal components of the receptor data (‘receptor space’). Brain regions clustered into rough anatomo-functional groups in receptor space. C) The receptor fingerprints of three areas at different points along the first principal component (i.e. the primary receptor gradient). The density of most receptors increases along the gradient, from are V1 to 3al and again to 12o. D) The first principal component closely follows the total receptor density per neuron.

### The density of receptors per neuron increases along the principal receptor gradient

Receptor fingerprints allow for visualization of the unique pattern of receptors expressed in each cortical area (Geyer et al., 1998; Zilles et al., 2002a). Cortical area V1 (visual cortex), 3al (somatosensory cortex) and 12o (orbitofrontal cortex) lie near the bottom, middle and top of the principal receptor gradient, while occupying similar positions along the secondary gradient. Visualizing the receptor fingerprints of these areas allows us to visualize changes in receptor expression along the principal receptor gradient. The most striking change along the principal receptor gradient was a general increase in the size of the receptor fingerprint, which is indicative of an increase in receptor density per neuron across almost all receptors (Fig 2C). Neurons near the top of the gradient contain on average a 3-4 times higher receptor density than those near the bottom (Fig 2D). The gradient closely tracked total receptors per neuron across brain areas (Fig 2D, r = 0.9953, p = 4x10^-110^). We next investigated how the principal receptor gradient relates to other known macroscopic gradients of anatomical organization.

### The principal receptor gradient increases along the cortical hierarchy

The cortical hierarchy describes an ordering in the cortex from areas that process basic sensory features of stimuli, low in the hierarchy, to areas that receive highly processed information, high in the hierarchy. Feedforward connections (from low to high areas in the hierarchy) tend to emerge from superficial cortical layers, while feedback connections (from high to low areas) tend to emerge from deep cortical layers (Markov et al., 2014b). Based on this knowledge, we recently estimated the cortical hierarchy of 40 cortical areas (Froudist-Walsh et al., 2020) using laminar retrograde tracing data (Felleman and Van Essen, 1991; Markov et al., 2014b). Here, we found a strong positive correlation between the principal receptor gradient and cortical hierarchy (r= 0.81, p = 8x10^-8^, all p-values Bonferroni corrected, Fig 3A). As the principal receptor gradient closely tracks the density of receptors per neuron, neurons lower in the cortical hierarchy (which process basic aspects of sensory stimuli) contain a relatively low number of receptors. In contrast, neurons near the top of the hierarchy (which receive multimodal information and contribute to complex cognitive functions) are endowed with a higher density of receptors, which may enable them to act with greater flexibility.

**Figure 3.**
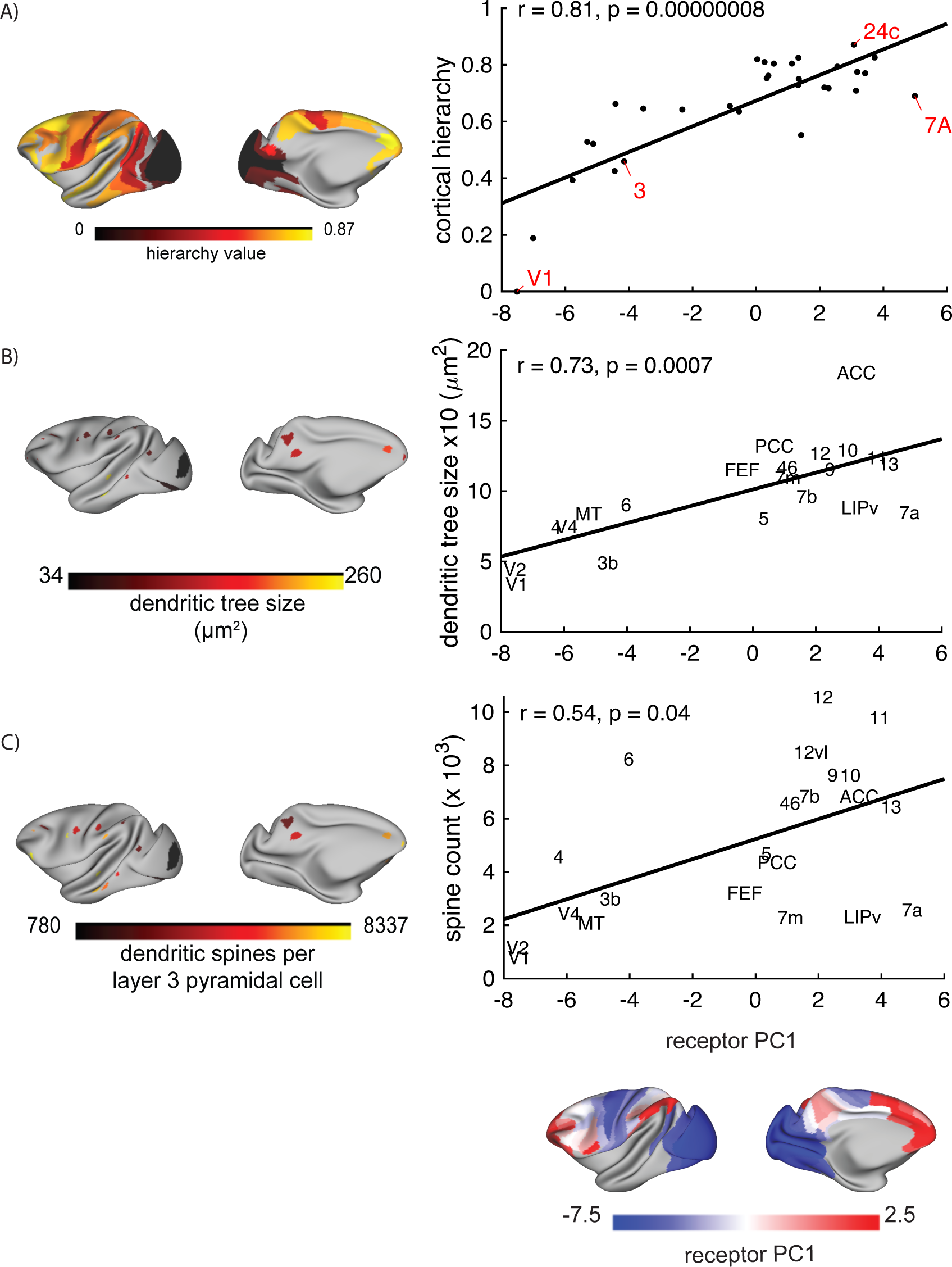
The anatomical foundations of the principal receptor gradient. A) The cortical hierarchy was estimated based on laminar connectivity data. There was a strong positive correlation between the principal receptor gradient and the cortical hierarchy. B and C) Dendritic tree size and spine count data were taken from a series of papers by Elston and colleagues (see text for references) and mapped to the common cortical space. Dendritic tree size and spines were both positively correlated with the principal receptor gradient.

### The size of the dendritic tree and number of dendritic spines increases along the principal receptor gradient

The striking increase in receptor density per neuron along the cortical hierarchy may require the emergence of a special cellular anatomy capable of housing such a high number of receptors. As pyramidal cells receive the vast majority of their synaptic contacts on the dendritic tree (Megias et al., 2001; Spruston, 2008), we investigated whether dendritic properties of neurons varied along the principal receptor gradient. Elston and colleagues measured the size of the dendritic tree and the number of dendritic spines on layer 3 pyramidal cells across dozens of areas of macaque cortex. Based on detailed descriptions and illustrations of the location of injections in those papers (Elston, 2001, 2000, 2000; Elston et al., 2011a, 2011a, 2011b, 2010, 2009, 2005, 1999; Elston and Rockland, 2002; Elston and Rosa, 1998a, 1997, 1998b), we identified the macroanatomic locations for the dendritic analyses on the Yerkes19 cortical surface. This revealed a positive correlation between the principal receptor gradient and dendritic tree size (r = 0.73, p = 0.0007; Fig 3B), and between the principal receptor gradient and the number of dendritic spines per neuron (r = 0.54, p = 0.04, Fig 3C). Thus, neurons that are higher up the principal receptor gradient contain larger dendritic trees, with more spines. This likely provides the neural real estate required to house a greater number of synaptic connections and receptors.

### Cortical areas and layers with high myelin content have low receptor density

Myelin has an inhibitory effect on plasticity (McGee et al., 2005), and part of the role of myelin in development may be to stop the creation of unwanted connections (Braitenberg, 1962). The ratio of T1-weighted to T2-weighted signal is strongly correlated with levels of myelination in the cortical grey matter (Glasser and Essen, 2011). We found a strong negative correlation between T1w/T2w ratio across the macaque cortex (obtained from Donahue et al., 2016) and the principal receptor gradient (Fig 4A, r = -0.72, p = 7x10^-19^). This suggests that neurons in cortical areas with a high myelin content express less neurotransmitter receptors.

**Figure 4.**
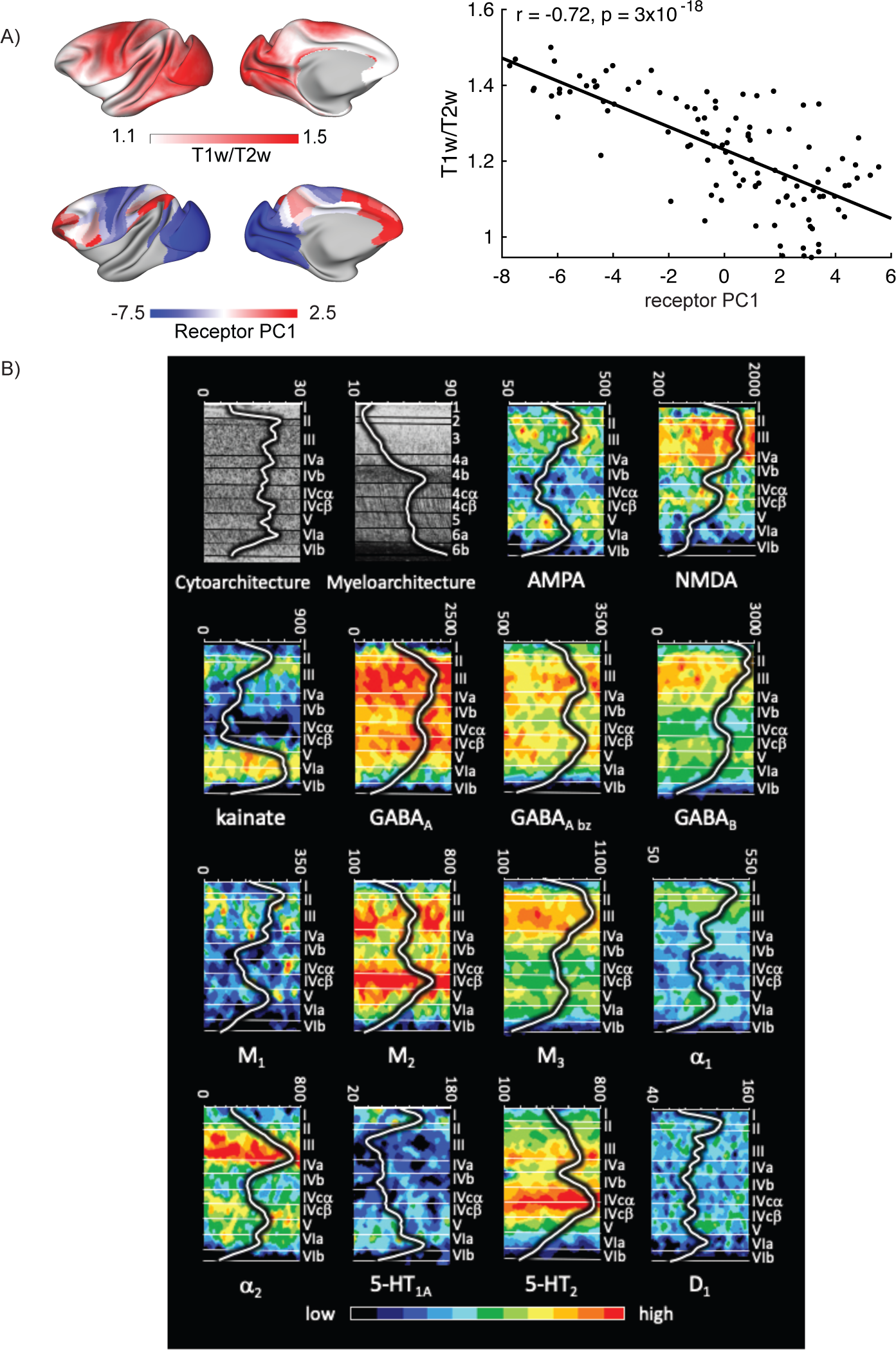
An inverse relationship between cortical myelin and receptor density. A) Cortical T1w/T2w ratio, a proposed marker for myelin content, is strongly negatively correlated with the principal receptor gradient across 109 cortical areas. B) Receptor-, cyto-, and myeloarchitecture of the macaque primary visual cortex (V1). The Grey Level Index, which represents a measure of the volume fraction of cell bodies, myelin density and receptor concentration (in fmol/mg protein) throughout the cortical depth is provided by the profile curve overlaid onto each section. Note that the scale has been optimized for each profile to provide the best visualization of changes in receptor densities throughout the cortical ribbon. Roman and Arabic numerals indicate cyto- and myeloarchitectonic layers, respectively. Positions of cytoarchitectonic layers were transferred to the neighboring receptor images.

There is a lack of theoretical understanding of the relationship between the T1w/T2w contrast and cortical myelin, which may break down under certain conditions, such as pathology (MacKay and Laule, 2016). To further probe the relationship between cortical myelin and receptor density, we compared the densities of all 14 receptors across cortical layers in V1 to the pattern of laminar myelination and cell density (Fig 4B). The cell density grey level index (GLI, which represents the volume fraction of cell bodies) was similar across all layers except for low densities in layers I and VIb. Myelin was expressed most strongly in layers IVb and VIb, and generally to a higher degree in infragranular than supragranular layers. Most receptors had highest densities in layers II and III, followed by layer V. Some receptors, including NMDA, GABA_A_, GABA_B_, M_2_ and 5HT_2_ also had high densities in layer IVc. Most receptor densities were low in layers I, VI and IVb. Thus, the receptor density pattern is opposite to the myelin density pattern across layers in area V1, particularly when accounting for differences in GLI levels across layers. Thus the receptor gradient, and receptor expression more generally, may be constrained by cortical myelination.

### The principal receptor gradient is significantly correlated with the two principal functional connectivity gradients

We next asked whether the principal receptor gradient may shape *in-vivo* interactions between cortical areas. A principal gradient of functional connectivity has previously been described in human cortex based on fMRI data (Margulies et al., 2016). Recently, Xu and colleagues identified shared gradients of functional connectivity across human and macaque cortex. This relied on quantifying the similarity of connectivity of each cortical point to each of 27 homologous cortical areas (Methods). We observed significant correlations between the principal receptor gradient and both the first (Fig 5A, r = 0.53, p = 4x10^-9^) and second (Fig 5B, r = 0.55, p = 4x10^-10^) functional connectivity gradients in macaque cortex.

**Figure 5.**
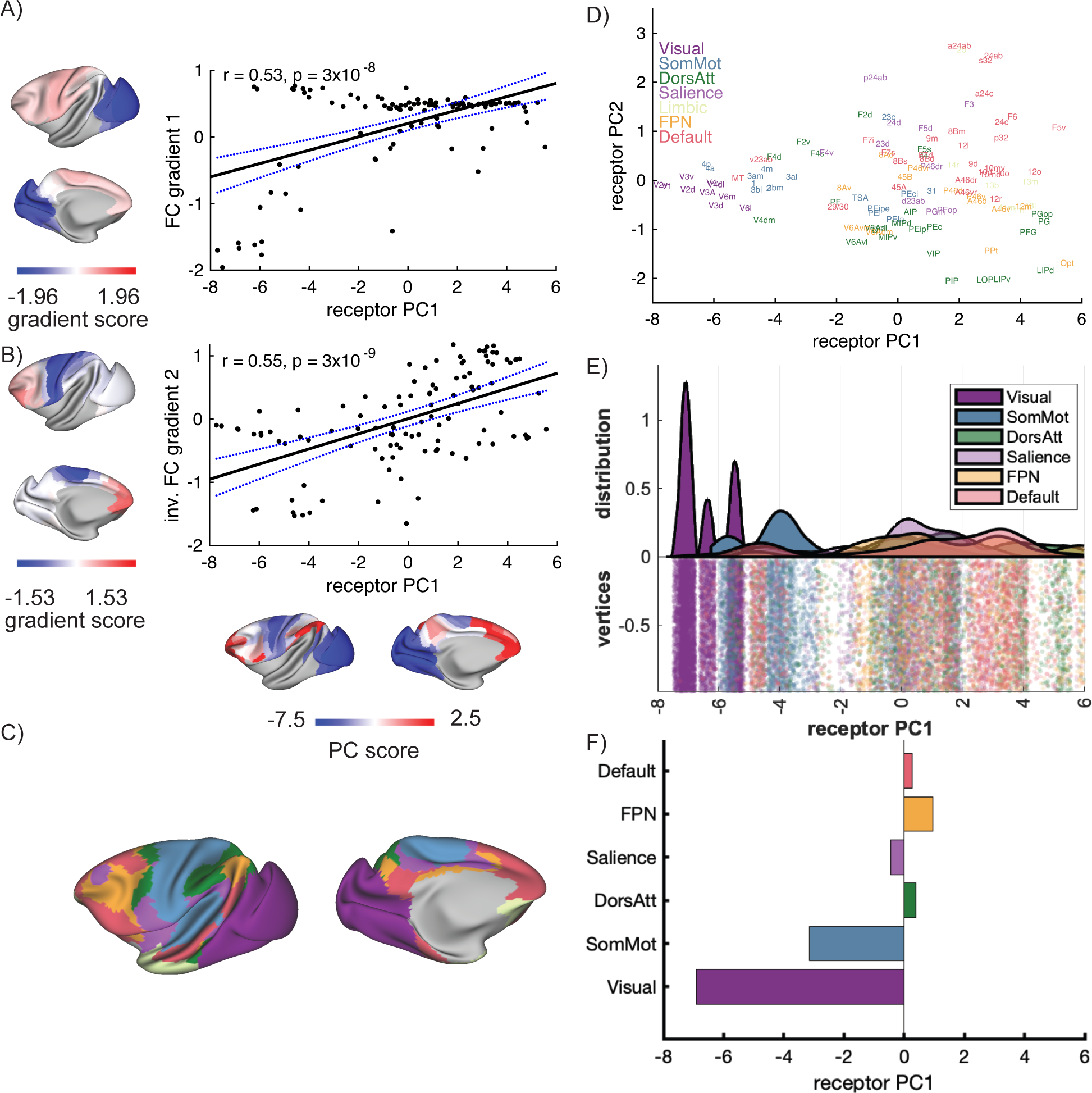
The principal receptor gradient underlies the principal gradients of in-vivo functional connectivity. A and B) The first and second gradients of functional connectivity across macaque cortex were calculated in (Xu et al., 2020). Both gradients were significantly correlated with the principal receptor gradient. C) The canonical cognitive networks of Yeo, Krienen and colleagues (Yeo et al., 2011) mapped to the macaque cortex by Xu and colleagues using cross-species functional alignment (Xu et al., 2020). D) The cognitive networks mapped into receptor space. E) The distribution of receptor PC1 scores for vertices in each cognitive network shown on a raincloud plot (Allen et al., 2019). The ‘limbic’ network was excluded, due to a lack of receptor data. F) The mean principal component score of the principal receptor gradient within each cognitive network.

The two functional connectivity gradients appeared to reflect different aspects of the primary receptor gradient, with the first connectivity gradient ranging from early visual cortex to frontal and parietal areas, and the second ranging from primary somatosensory and motor areas to prefrontal cortex.

The shared functional connectivity gradients were used by Xu and colleagues to identify corresponding points in human and macaque cortex that best preserved global cortical connectivity patterns (Xu et al., 2020). This provided a cross-species functional alignment, which was used to align the canonical seven cognitive networks of Yeo, Krienen and colleagues (Yeo et al., 2011) from the human cortex to the macaque cortex (Xu et al., 2020) (Fig 5C). We used this cross-species functional alignment to identify the receptor gradient expression across cognitive networks. The overlap of each area of the Julich Macaque Brain Atlas with the seven cognitive networks is quantified in Supplementary Table 1. We excluded the ‘limbic’ network due to the lack of vertices with receptor data, and focused on the remaining six networks. We found that the principal receptor gradient clearly separated the sensory (visual and somatomotor) networks, which had low receptor gradient scores, from the higher cognitive networks (dorsal attention, salience, frontoparietal and default mode), which had relatively higher gradient scores (Fig 5D-F). Almost all areas of the visual and somatosensory networks had negative gradient scores, while areas in the higher cognitive networks encompassed a range of positive and negative values (Fig 5D,E).

Taken in conjunction with findings above, this suggests that higher cognitive networks contain neurons with a relatively high density of receptors, which may contribute to their flexibility of function.

### The secondary receptor gradient reflects variation in serotonin 5-HT_1A_ receptors between higher order cortical areas

The secondary receptor gradient separated higher order cortical areas (Fig 6A). Parietal areas such as LIPv lay at one end of the gradient, while cingulate areas such as 24ab and 25 lay at the other (Fig 6A). Visualization of the receptor fingerprints for representative areas revealed a striking difference in the serotonin 5-HT_1A_ receptor density, as well as differences in the GABA_A_ density and AMPA- kainate/NMDA ratio between areas at opposite ends of the secondary receptor gradient (Fig 6B). This was confirmed by correlations with secondary receptor gradient (Fig 6C) and with visualization of the coefficients of the secondary principal component (Fig S2). Areas at the top of the secondary receptor gradient, located in the subcallosal and anterior cingulate cortex, had a particularly high 5-HT_1A_ receptor density. There was no strong relationship between serotonin 5-HT_2_ expression and the secondary receptor gradient (Supp. Fig. 3). The 5-HT_1A_ is the principal serotonin receptor that has a predominantly hyperpolarizing effect on neurons in the cortex (Celada et al., 2013).

**Figure 6.**
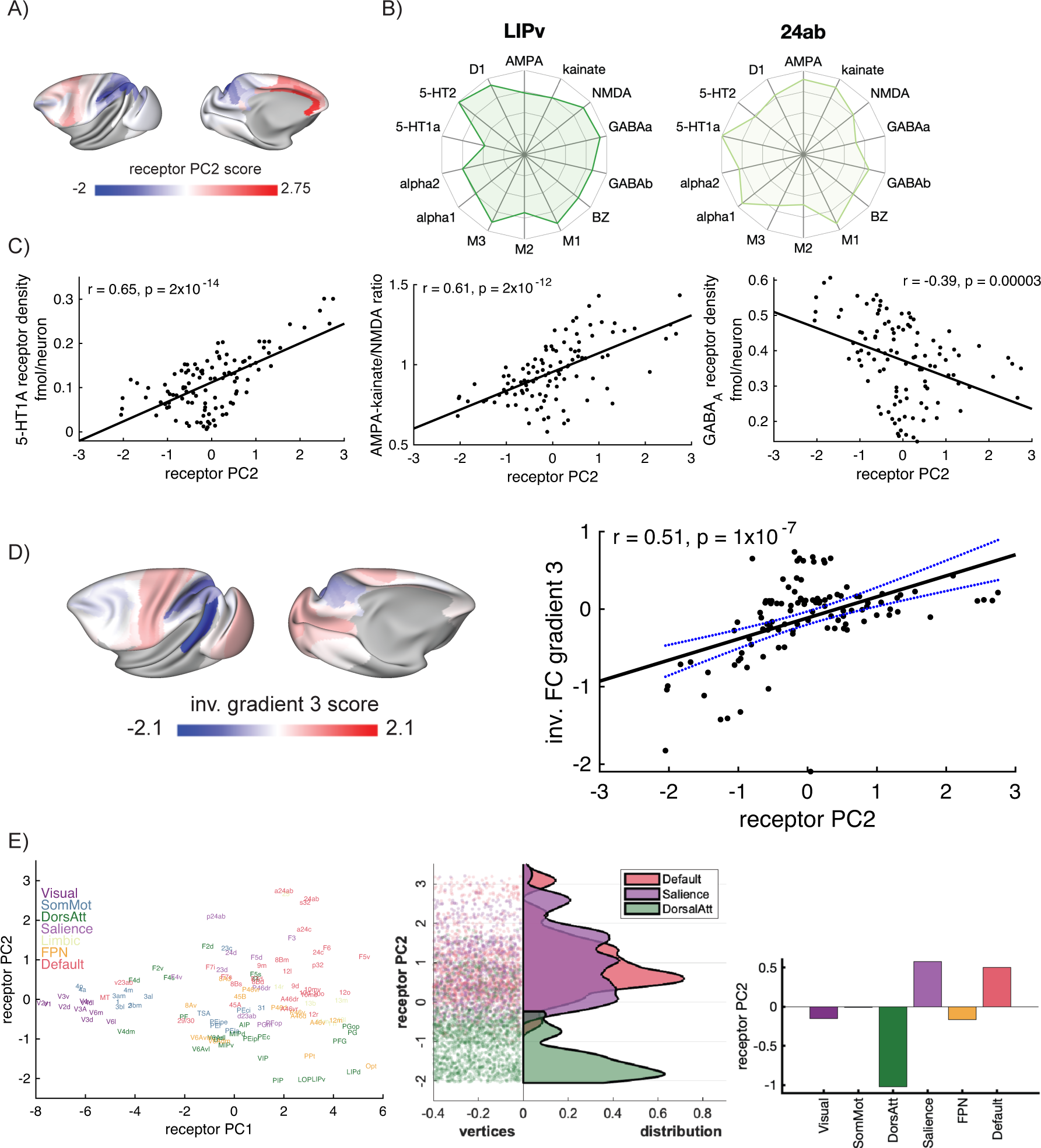
The secondary receptor gradient tracks differences in serotonin receptor densities between higher cognitive areas of cortex. A) Receptor PC2 (the secondary receptor gradient). B) Receptor fingerprints of areas 24ab and LIPv, which occupy opposing positions along the secondary gradient. The total receptor density per neuron is similar between the two areas, but there is an obvious difference in 5-HT_1A_ receptor density per neuron. C) Receptor PC2 was positively correlated with 5-HT_1A_ receptor density and the ratio of AMPA and kainate to NMDA receptors. Receptor PC2 was also negatively correlated with the GABA_A_ receptor density.D) Left. The third functional connectivity gradient. Right. The secondary receptor gradient was strongly correlated with the third functional connectivity gradient. E) The secondary receptor gradient separates the dorsal attention network from the default mode and salience networks. Left, cognitive networks in receptor space. Middle, distribution of receptor PC2 scores within the default mode, salience and dorsal attention networks. Right. Mean receptor PC2 scores for each cognitive network.

Interestingly, major depressive disorder is associated with reduced serotonin signaling, and deep brain stimulation of the subcallosal white matter has been shown to be an effective treatment of depression in humans (Mayberg et al., 2005). Therefore, our finding provides a potential anatomical and molecular explanation for why selective serotonin reuptake inhibitors (SSRIs) reduce glucose metabolism in the subgenual cingulate in human patients with depression.

### The secondary receptor gradient separates the dorsal attention network from the default mode network and salience network

We then investigated whether the secondary receptor gradient also corresponded to gradients of functional connectivity. Indeed, the secondary receptor gradient was significantly correlated with the third functional connectivity gradient (Fig 6D), with both gradients anchored in the lateral parietal cortex and precuneus. In contrast to the principal receptor gradient, which aligns with the cortical hierarchy, the secondary receptor gradient separates parietal and cingulate areas, which contribute to higher cognitive functions, and could thus facilitate switching between higher cognitive states in the brain.

We then analyzed the secondary receptor gradient score of each cognitive network (Fig 6E). There was a strong negative weighting in the dorsal attention network, and strong positive weighting in regions of the default mode network. These two higher cognitive networks typically are anticorrelated (Chai et al., 2012; Fox et al., 2005; Kelly et al., 2008; Yeo et al., 2015). Additionally, there was a strong positive loading in the salience network, which has been proposed to act as a ‘switch’ between default mode network activity and task-engaging frontoparietal network activity (Menon and Uddin, 2010). Thus, the secondary receptor gradient, and in particular the 5-HT_1A_ receptor expression pattern may capture a mechanism by which the cortex can switch between higher cognitive states dominated by the dorsal attention network or the default mode network.

### Macaque 5-HT_1A_ receptor and human HTR1A gene expression peak in the default mode and salience networks

We then investigated the correspondence between human HTR1A gene expression and macaque 5-HT_1A_ receptor expression. The HTR1A gene codes for the 5-HT_1A_ receptor. To do this we examined gene expression from the Allen Human Brain Atlas. This contains mRNA expression from hundreds of anatomical locations, with the left cortical hemisphere heavily sampled across 6 individual brains (Hawrylycz et al., 2012). We mapped gene expression onto a comprehensive 180 area multimodal parcellation of the human cortex (Glasser et al., 2016). HTR1A gene expression peaked in anterior/medial temporal cortex, insula, subcallosal/anterior cingulate and areas of postero-medial cortex (posterior cingulate/precuneus) (Fig 7A). We reverse-translated this map to the macaque cortex using cross-species functional alignment (Xu et al., 2020) (Fig 7A, Methods). There was a strong positive correlation between human HTR1A gene and macaque 5-HT_1A_ receptor expression (r = 0.66, p = 6x10^-15^, Fig 7B).

**Figure 7.**
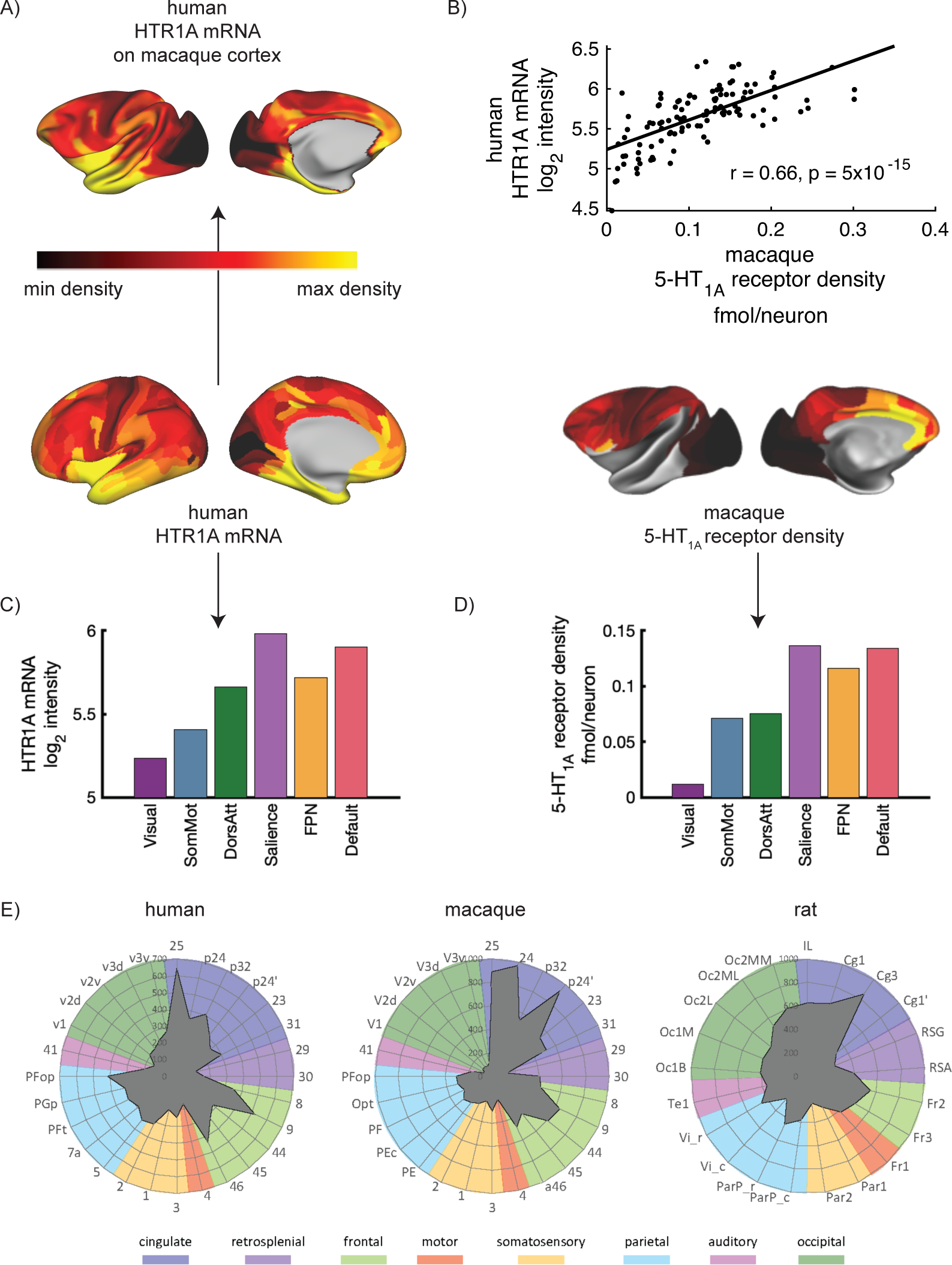
Serotonin 5-HT_1A_ receptor expression across the human, macaque and rat cortex. A) We mapped human HTR1A gene expression data (Hawrylycz et al., 2012) to the human cortex and then to the macaque cortex using cross-species functional alignment. B) Human gene expression and macaque receptor expression for the 5-HT_1A_ receptor were positively correlated. C and D) Human HTR1A gene expression and macaque 5-HT_1A_ receptor expression are expressed similarly across cognitive networks, peaking in the Default Mode and Salience networks. E) The density of 5-HT_1A_ receptors (in fmol/mg protein) across multiple areas of human, macaque and rat cortex. 5-HT_1A_ receptor density is high in subcallosal or anterior cingulate in all species, although the area with peak density shifts slightly across species.

Both human HTR1A gene and macaque 5-HT_1A_ receptors were expressed similarly across cognitive networks, with the order of gene/receptor expression preserved (from lowest to highest expression: visual, somatomotor, dorsal attention, frontoparietal, default, salience; Fig 7C,D). Note that this expression is largely captured by the first two receptor gradients. The low expression in visual and somatomotor networks is captured by the principal receptor gradient. The ordering of the higher cognitive networks is captures by the secondary receptor gradient.

### 5−HT_1A_ receptor expression across cortex is largely conserved across species

We then assessed the pattern of 5-HT_1A_ receptor expression across cortex in the human, macaque and rat using *in-vitro* receptor autoradiography (Fig 7E). In the human, 5-HT_1A_ receptor expression peaked in area 25 (subcallosal cingulate), with high density in other anterior cingulate and frontal regions and low density in motor and visual cortex. This general pattern was also true in macaque and rat cortex. However, the peak of 5-HT_1A_ receptor density appears to shift slightly from area 25 to neighboring parts of anterior cingulate in the macaque, with a similar shift even more apparent in the rat. In the rat, the gradient of 5-HT_1A_ receptor density across cortex was generally similar to that seen in the macaque and human, but was somewhat flatter.

## Discussion

In this work, we measured expression of 14 receptor types per neuron across 109 regions of macaque cortex, and described their general organizational principles. We discovered a principal gradient of cortical receptor expression along the cortical hierarchy that accounted for about 80% of variance in the entire receptor dataset. This ‘principal receptor gradient’ tracked the increased expression of all receptor types from early sensory cortex to regions of cortex contributing to higher cognitive processes. We identified a plausible neural architecture to house this increased receptor density. Brain regions with more receptors per neuron contained pyramidal cells with larger dendritic trees, more dendritic spines, and displayed a lower T1w/T2w ratio in MRI scans, indicative of reduced grey matter myelin. The inverse relationship between receptor density and myelin was confirmed across layers in V1. The patterns of receptor expression were strongly correlated with *in-vivo* gradients of functional connectivity, suggesting a neurochemical basis for large-scale cortical connectivity patterns. The secondary receptor gradient was largely driven by expression of the serotonin 5-HT_1A_ receptor. The pattern of macaque 5-HT_1A_ receptor expression across cortex was very similar to human 5HT_1A_ receptor and gene expression, and to a lesser degree to rat 5HT_1A_ receptor expression. The second gradient segregated the dorsal attention from the default mode network and salience network. This suggests the possibility of a serotonergic basis for a switch between external and internal modes of attention.

In recent years, the description of cortical organization in terms of gradients of gradually changing properties (Braitenberg, 1962; Sanides, 1962) has returned to fashion (Wang, 2020). Gradients of connectivity (Margulies et al., 2016; Xu et al., 2020), cell-type densities (Kim et al., 2017), receptor expression (Goulas et al., 2021) and gene expression (Burt et al., 2018; Fulcher et al., 2019) have been described. Many of these properties vary along an axis that aligns with the cortical hierarchy, which can be estimated based on patterns of laminar connectivity (Felleman and Van Essen, 1991; Froudist-Walsh et al., 2020; Markov et al., 2014b; Theodoni et al., 2020). One of the mysteries of such organization is how gradual changes in anatomical properties can lead to the emergence of strikingly different functions. Here we described a principal receptor gradient in macaque cortex that increased along the hierarchy. Along the principal receptor gradient, we show that neurons that are further up the hierarchy express on average 4 times higher densities of neurotransmitter receptors than neurons at the lowest end of the hierarchy. This includes not only the ubiquitous excitatory glutamate and inhibitory GABA receptors, but also 8 types of neuromodulatory receptors. We found that neurons with more receptors also contain much larger dendritic trees and more spines, are thus anatomically equipped to integrate information from a larger number of sources. This coincides with the increased neuronal timescales with which neurons in higher areas integrate information (Chaudhuri et al., 2015; Gao et al., 2020; Hasson et al., 2008; Murray et al., 2014; Raut et al., 2020). This extra receptor capacity may endow neurons in higher cortical areas with the ability to respond flexibly when faced with changing demands. In contrast, the receptor-light neurons of early sensory cortex may receive relatively little modulation, which may ensure that sensory stimuli are processed reliably in different contexts.

The second receptor gradient was dominated by serotonin 5-HT_1A_ receptor expression. This peaked in the cingulate cortex, with high expression in the subcallosal cingulate, which is a principal target of deep brain stimulation for treating depression. The extremely high expression of 5-HT_1A_ receptors in this same brain regions provides a plausible explanation for why deep brain stimulation of the subcallosal cingulate and selective serotonin reuptake inhibitors (SSRIs) have almost identical effects on cerebral blood flow (Mayberg et al., 2005, 2000). SSRIs increase the activation of 5-HT_1A_ receptors, which are most prominently expressed in subcallosal and anterior cingulate (Fig 7), where they act to reduce neural activity and counteract the increased glucose metabolism seen in patients with depression in this area (Mayberg, 2003, 1997). Although gene expression is not always a good predictor of receptor expression (Arnatkeviciute et al., 2019; Schwanhäusser et al., 2011), a previous study using PET found that gene and receptor expression are closely aligned for the 5-HT_1A_ receptor in humans (Beliveau et al., 2017). Here, using exquisite resolution of *in-vitro* autoradiography, we found that 5-HT_1A_ receptor density in the macaque was very similar to the HTR1A gene expression and receptor expression in humans (Fig 7). The pattern of 5-HT_1A_ receptor expression in the rat also peaked in the cingulate cortex, however the gradient of expression was flatter in the rat than in the macaque or human brain. Notably, the laminar receptor expression pattern in the rat differs considerably from that observed in human or macaque cortex (Zilles and Palomero-Gallagher, 2017a), and serotonin receptors are expressed on different cell types between the mouse and human (Hodge et al., 2019). The differences between the primate and rodent receptor expression across brain areas, layers and cell types suggest caution when designing and interpreting rodent models of depression. In contrast, the closely overlapping serotonin receptor anatomy between macaque monkey and human reported here, suggests that, the macaque may be a promising animal model for depression.

In contrast to the principal receptor gradient, which separates early sensory from higher cognitive areas, the second receptor gradient segregated higher cognitive networks. In particular, the second gradient separated the dorsal attention network from the default mode network and salience network. Association cortex is often divided into four distributed cognitive networks (dorsal attention, salience, frontoparietal, default), with a fifth higher cognitive network, the ‘limbic network’ often included within the default network (Uddin et al., 2019). These networks each occupy parts of the frontal, parietal and temporal lobes, and in several patches of cortex appear in a consistent order along the cortical surface (Margulies et al., 2016). The dorsal attention network usually lies closest to sensory areas, and is highly active when attention must be focused on external stimuli (Corbetta and Shulman, 2002). The default mode network lies furthest away from sensory areas, and is deactivated during similar tasks, but highly active during tasks that require a disconnection of attention from the external world, such as autobiographical memory, or imagination (Andrews-Hanna et al., 2014; Buckner et al., 2008; Raichle, 2015; Shulman et al., 1997; Spreng and Grady, 2009). Activity in these two networks is often anticorrelated (Chai et al., 2012; Fox et al., 2005; Kelly et al., 2008; Yeo et al., 2015), in line with their opposing roles in cognition. The frontoparietal network (known by several other names, including the multiple demand system, the cognitive control network and the central executive network (Uddin et al., 2019)), lies between these two networks and may dynamically couple with either, depending on task demands (Dixon et al., 2018; Spreng et al., 2010). In line with this intermediate role, the frontoparietal network lay between the dorsal attention and default mode networks along the secondary receptor gradient.

Previous models have suggested that the antagonism between the dorsal attention network and the default mode network may result from direct long-range excitatory connections that target inhibitory neurons in the opposing network (Anticevic et al., 2012). We propose that the cognitive networks may represent distinct attractor states of the large-scale system, which can be seen as relatively stable patterns of spatiotemporal activity. Based on our findings, release of serotonin would engage inhibitory 5-HT_1A_ receptors principally in the default-mode network, and the salience network. Previous evidence suggests that the salience network acts as a switch from the default mode network to frontoparietal networks focused on external attention (Menon, 2011; Menon and Uddin, 2010). Serotonin neuron activity closely resembles a “surprise” signal (Matias et al., 2017), and surprising stimuli activate the salience network (Menon, 2015), which may engage the dorsal attention network to focus attention externally towards such stimuli. The distribution of 5-HT_1A_ receptors on excitatory and inhibitory cell types (Puig et al., 2010; Puig and Gulledge, 2011; Xiang and Prince, 2003) in the default mode and salience networks may determine whether serotonin release has a similar effect on both networks. The 5-HT_1A_ receptor is thought to dominate cortical serotonin processing under normal conditions, due to its high affinity for serotonin compared to 5-HT_2_ receptors (Hoyer et al., 1986a, 1986b, 1985). In contrast, the 5-HT_2_ system is engaged by massive serotonin release under extreme conditions (Carhart-Harris and Nutt, 2017), when attention needs to be rapidly shifted to external events. Accordingly, the excitatory effects of the 5-HT_2_ receptors may complement the 5-HT_1A_ effects by preferentially exciting the dorsal attention and frontoparietal networks (Fig S3). This is compatible with recent findings that genes for neuromodulatory receptors are expressed at cortical locations that may affect the flow of brain states over time (Shine et al., 2019), and suggests a specific mechanism for shifting activity between cardinal cognitive networks.

The receptor architecture of human cortex has recently been described (Zilles and Palomero-Gallagher, 2017b), with dimensionality-reduction techniques finding a gradient comparable to the principal receptor gradient we present here (Goulas et al., 2021; Zilles and Palomero-Gallagher, 2017b). However, there are a few key differences with the present study. First, we study macaque cortex, which allows for comparison with gold-standard invasive anatomy, imaging and physiology data. Second, analyzing the macaque cortex allowed us to combine receptor autoradiography data with neuron density across cortex (Collins et al., 2010). This enabled us to estimate the density of 14 types of receptor per neuron, which is important to understand the functional significance of changes in receptor densities. One clear result that emerges from processing the data in this manner is that several receptors that have high density (in the raw data) in V1 do so principally because of the remarkably high neuron density in that area. Third, we were able to bring multiple invasive anatomical and *in-vivo* functional datasets into a common space, which enabled us to identify the anatomical structure underpinning the receptor gradients, as well as discover the potential significance of such gradients in guiding functional communication in the cortex.

Recent developments in large-scale recording techniques have highlighted the distributed nature of cognitive functions. Despite these advances, our theoretical understanding of how distinct cognitive functions emerge across different areas of the cortex is limited. The large-scale receptor data presented here can provide an anatomical basis for future large-scale models of neuromodulation of brain connectivity (Deco et al., 2018), activity and cognitive function (Cano-Colino et al., 2014; Froudist-Walsh et al., 2020) and for understanding the emergence of flexible higher cognition along the cortical hierarchy.

## Methods

### In-Vitro Receptor Autoradiography of the Macaque Cortex

We analysed the brains of three adult male *Macaca fascicularis* specimens (between 6 and 8 years old; body weight between 5.2 and 6.6 kg) obtained from Covance Preclinical Services GmbH, Münster, where they were used as control animals for pharmaceutical studies performed in compliance with legal requirements. All experimental protocols were in accordance with the guidelines of the European laws for the care and use of animals for scientific purposes.

Animals were sacrificed by means of an intravenous lethal dose of sodium pentobarbital. Brains were removed immediately from the skull, and brain stem and cerebellum were dissected off in close proximity to the cerebral peduncles.

Hemispheres were separated and then cut into a rostral and a caudal block by a cut in the coronal plane of sectioning within the central sulcus. These blocks were frozen in isopentane at -40°C to -50°C, and then stored in airtight plastic bags at -70°C. Each block was serially sectioned in the coronal plane (section thickness 20 µm) using a cryostat microtome (CM 3050, Leica, Germany).

Sections were thaw-mounted on gelatine-coated slides, freeze-dried overnight and processed for visualization of receptors, cell bodies (Merker, 1983) or myelin (Gallyas, 1979).

Quantitative *in vitro* receptor autoradiography was applied to label 14 receptors from different neurotransmitter systems according to previously published protocols (Palomero-Gallagher and Zilles, 2018b; Zilles et al., 2002b; Supplementary Table 1), and encompassing a preincubation to rehydrate sections, a main incubation with a tritiated ligand in the presence of or without a non-labeled displacer, and a final rinsing step to terminate binding. Incubation with the displacer enabled to determine the proportion of displaceable, non-specific binding, which was less than 5% of the total binding. Thus, total binding is considered to be equivalent of specific binding. Sections were dried in a cold stream of air, exposed together with plastic scales of known radioactivity against tritium-sensitive films (Hyperfilm, Amersham) for 4-18 weeks depending on the receptor type.

Ensuing autoradiographs were processed by densitometry with a video-based image analysing technique (Palomero-Gallagher and Zilles, 2018b; Zilles et al., 2002b). For details of the method, please see (Palomero-Gallagher and Zilles, 2018b; Zilles et al., 2002b). The mean receptor density for each area was determined by density profiles extracted vertical to the cortical surface over a series of 3–5 sections per receptor type and animal using Matlab-based in house software (Palomero-Gallagher and Zilles, 2018b). Identification of cortical areas and layers was carried out by comparison with the adjacent cell-body stained histological sections. Finally, autoradiographs were pseudo-colour coded by linear contrast enhancement and assignment of equally spaced density ranges to a spectral arrangement of eleven colours for visualization purposes.

### Creation of surface representation of cyto- and receptor-architectonic atlas and receptor data

One-hundred-and-nine cortical areas were defined based on their receptor- and cyto-architecture, as described in (Impieri et al., 2019; Niu et al., 2021, 2020; Rapan et al., 2020) and upcoming publications on the prefrontal cortex, cingulate and occipital lobe. We call this parcellation the Julich Macaque Brain Atlas. The location and extent of the cortical areas were delineated in the 3D space of the Yerkes19 surface (Donahue et al., 2016) by LR, MN and NPG using the connectome workbench software (https://www.humanconnectome.org/software/connectome-workbench) by carefully aligning boundaries to macroanatomical landmarks identified using the cytoarchitecture. The location of all regions on the Yerkes19 surface were independently checked and verified by MN, SFW, LR and NPG. 3D reconstruction of the hemisphere was obtained using the Connectome Workbench software. Additionally, the mean receptor densities of all 14 receptor types have been projected onto the corresponding area on the Yerkes19 surface for visualization.

### Creation of surface representation of regions for neural density data

Collins and colleagues measured the neural density across the macaque cortex (Collins et al., 2010) using the isotropic fractionator method (Herculano-Houzel and Lent, 2005). In that paper, the cortex is presented as a flat-map divided into sections in their Figure 2 and S3. We used these maps, along with several sulcal, gyral and areal landmarks provided in their Figure 2 to estimate the location of each cortical section (i.e. the Vanderbilt sections) on the Yerkes19 surface. This was performed by SFW and independently verified by LR, MN and NPG. This allowed us to estimate the neural density in each of the 109 areas of the Julich Macaque Brain Atlas. For each area in the Julich Macaque Brain Atlas, we took a weighted average of the neural density in each of the overlapping Vanderbilt sections, weighted by the number of vertices of overlap. The neuron density data was originally in units of neurons per gram, and the receptor density data in fmol/mg protein. To estimate the receptor density in fmol per neuron, we used the previously reported figure that 8% of brain tissue is protein (McIlwain and Bachelard, 1972). This amounts to multiplying by a constant, and does not affect the calculation of the gradients via principal components analysis or the correlations with other maps.

### Receptor gradients

To identify the receptor gradients, we z-scored the receptors-per-neuron data and performed a principal components analysis. Z-scoring ensured that high density receptors would not dominate the principal components.

### Cortical hierarchy and retrograde tracing data

The cortical connectivity data were obtained from Henry Kennedy (Lyon, France) and are available at core-nets.org. The retrograde tracing data was obtained following injections into 40 cortical regions using consistent methods in the same laboratory. This ensures high quality and consistency of weighted and directed connectivity data (Kennedy et al., 2013). This database is regularly updated with new connectivity data (Markov et al., 2014a). We recently estimated the cortical hierarchy using this data, based on the laminar patterns of connections (Froudist- Walsh et al., 2020). For details of the method, please see (Froudist-Walsh et al., 2020; Markov et al., 2014b). The parcellation for this connectivity data (the Lyon Macaque Brain Atlas; known in BALSA as the M132 atlas) has previously been made available on the Yerkes19 surface (Donahue et al., 2016). We used this to fill in hierarchy values on the surface. We then estimated the receptor gradient values within each area of the Lyon Macaque Brain Atlas, weighted by the extent of overlap with each area of the Julich Macaque Brain Atlas.

### Creation of surface representation of dendritic data

Elston and colleagues measured the dendritic tree length and spine density of layer III pyramidal neurons in a series of studies. In order to compare this data with the receptor data, we first mapped the injected regions as described by Elston and colleagues onto the Yerkes19 template. Borders for injection sites in the series of papers by Elston and colleagues were drawn on the Yerkes19 template by SFW. Direct comparison with the hand-drawn maps was possible for areas V1, V2, MT, LIPv, 7a, V4, TEO, STP, IT, Ant. Cing., Post. Cing, TEpd, 12vl, A1, 3b, 4, 5, 6, 7b, 9, 13, 46, 7m (Elston, 2001; Elston et al., 2011a, 2010, 2009, 2005, 1999; Elston and Rockland, 2002; Elston and Rosa, 1998a, 1997). Direct comparison with the anatomical drawings was supplemented by additionally comparing the drawn locations with the anatomical references cited therein, and, where possible, atlases available within connectome workbench that were cited in the studies by Elston. Areas 10, 11 and 12 (Elston, 2000) were described with reference to (Preuss and Goldman-Rakic, 1991). The injection in area TEa, as described in (Elston et al., 2001) used the maps in (Seltzer and Pandya, 1978) for area definition. We used these maps to approximate the injection location. Area STP, injected in (Elston, 2001; Elston et al., 1999) was identified with the corresponding region STPp in the atlas of Felleman and Van Essen (Felleman and Van Essen, 1991). Area FEF was identified according to the description in (Elston and Rosa, 1998b), lying on the anterior bank of the medial aspect of the arcuate sulcus. All identified injection sites on the cortical surface were independently verified by MN, LR and NPG. The receptor principal component score was averaged within all vertices in each injection site in order to compare dendritic and receptor data.

### Cortical T1w/T2w data

The T1w/T2w data was acquired by (Donahue et al., 2016), and was downloaded from the BALSA neuroimaging website (Van Essen et al., 2017). To compare the T1w/T2w data with the receptor data, we simply averaged the T1w/T2w signal within each of the 109 areas of the Julich Macaque Brain Atlas.

### Functional connectivity data

Xu and colleagues recently identified matching functional connectivity gradients in human and macaque cortex (Xu et al., 2020). These matched gradients were used to develop a mapping between human and macaque cortex that best aligns points (vertices) in macaque and human cortex according to their global functional connectivity patterns (Xu et al., 2020). Here we used the human-to-macaque mapping developed in that manuscript. We provide some details of how that mapping was calculated.

Resting-state fMRI data from 19 macaque monkeys was collected with no contrast agent in Oxford (Noonan et al., 2014). These data were downloaded from the PRIME-DE database (Milham et al., 2018) and pre-processed using a Human Connectome Project-like pipeline (Xu et al., 2015, 2019, 2018). Similarly, minimally preprocessed human resting-state fMRI data from the Human Connectome Project was used (Glasser et al., 2013).

Gradients of functional connectivity were defined using the diffusion map method (Coifman and Lafon, 2006; Margulies et al., 2016). Diffusion maps rely on a measure of similarity between each cortical point. In previous studies, the similarity of functional connectivity patterns between cortical vertices was used to define the similarity matrix (Margulies et al., 2016). To compare macaque and human cortex, a cross-species joint-similarity matrix had to be used as input to the diffusion mapping.

The diagonal blocks of the joint-similarity matrix (corresponding to similarity of human to human cortical vertices, and monkey to monkey cortical vertices) were based on (cosine) similarity of functional connectivity patterns, as in previous studies (Margulies et al., 2016). To create a similarity measure between each point on the macaque cortex and each point on the human cortex, 27 landmark regions were used (related to the approach by (Mars et al., 2018)). These had previously been identified as homologous regions between human and macaque monkeys (Mars et al., 2011; Neubert et al., 2014; Sallet et al., 2013; Van Essen and Dierker, 2007). For each species, the similarity between the functional connectivity patterns of each vertex and each landmark were calculated (for a similar approach, see (Mars et al., 2018)). Then the human vertex to monkey vertex similarity could be calculated by assessing how their connectivity resembled that of the 27 landmark regions. The matrix containing this (cosine) similarity for each pair of human and monkey vertices, and its transpose, made up the off-diagonal squares of the joint-similarity matrix.

This cross-species joint-similarity matrix was used as input to the diffusion map. This produced shared cross-species gradients of functional connectivity. To create the point-to-point cross-species mapping, these functional connectivity gradients were used as input to multimodal surface matching (MSM) (Robinson et al., 2014), the method recently used to define a high-quality parcellation of human cortex based on multimodal imaging features (Glasser et al., 2016). Once this mapping is established, it can be used to transfer maps or parcellations across species. Xu et al demonstrated the accuracy of this method through accurate alignment of myelin maps across species. However, even for best-matching homologous points of cortex, there are cross-species differences in functional connectivity patterns. These cross-species functional connectivity differences are greater for associative than sensory regions of cortex (Xu et al., 2020).

The definition of seven cognitive networks in human cortex by Yeo, Krienen and colleagues has become a standard network definition in the field (Yeo et al., 2011). Some authors have since suggested that the limbic network in fact forms part of the default mode network, leaving six canonical cognitive networks (Uddin et al., 2019). Xu et al transferred the cognitive networks defined in (Yeo et al., 2011) from human to macaque using this cross-species functional alignment (Xu et al., 2020). We used this human-to-monkey mapping to identify the receptor expression across cognitive networks. Due sparse receptor mapping in the limbic network, we excluded this cognitive network from analysis.

### Human gene expression data

Human gene expression data was downloaded from the Allen Human Brain Atlas (Hawrylycz et al., 2012). We analyzed data from hundreds of microarray samples across the left cortical hemispheres of six donors (5 male, 1 female). We replicated the methods in (Burt et al., 2018) using in-house code in Matlab (github.com/seanfw/genemapper), with the following exceptions: 1) Instead of using the MNI coordinates supplied by the Allen Institute, we used the native-space sample coordinates, and performed a surface registration of the individual brains to the HCP group average surface. Surface registration is well suited to detect the sulcal patterns of postmortem brains, and relatively unaffected by nonlinear deformations to subcortical structures and white matter (such as squishing) that can affect postmortem brains. 2) Rather than mapping genes to brain areas for each subject and then averaging across subjects, we mapped genes to brain areas for all subjects together. This reduces the need to interpolate values for brain areas that contain no samples, but may be more vulnerable to individual differences in gene expression. Similar to (Arnatkeviciute et al., 2019), we found that the spatial patterns of gene expression were similar regardless of whether the gene expression data were normalized by z-scoring across samples or across genes. We extracted the HTR1A gene expression pattern, and mapped this to the macaque cortex using the cross-species functional alignment detailed above (Xu et al., 2020).

### Statistical analysis

Following mapping of all data to a common space, Pearson correlations were performed between the receptor principal components and each of the other data-types mentioned above. P-values were Bonferroni corrected based on the number of correlations between receptor gradients and structural or functional maps.

## Acknowledgments

This project was funded by (NIH/BMBF) CRCNS grant (nos. R01MH122024 and 01GQ1902) to NPG and XJW; NIH grant R01MH062349, ONR grant N00014 and James Simons foundation grant 543057SPI to XJW and the European Union’s Horizon 2020 Framework Programme for Research and Innovation under the Specific Grant Agreements 785907 (Human Brain Project SGA2) and 945539 (Human Brain Project SGA3) to KZ and NPG. We would also like to thank the investigative teams from Oxford (Sallet, Mars, Rushworth), Washington University (Donahue, Glasser, Van Essen and colleagues), the Allen Institute (Hawrylycz and colleagues), Vanderbilt (Collins, Kaas and colleagues), Queensland (Elston and colleagues) and Lyon (Kennedy and colleagues) for collecting, publishing and making available their original data.

## Author contributions

Conceptualization - SFW, NPG, XJW. Methodology - SFW, TX, DM, KZ, NPG, XJW. Software – SFW, TX.

Validation - SFW, MN, LR, NPG, XJW. Formal Analysis - SFW. Investigation - NPG, LR, MN, KZ. Resources - NPG, KZ, XJW.

Writing - original draft preparation - SFW. Writing - review and editing - all authors. Visualization – SFW, LR, NPG.

Supervision - NPG, KZ, XJW.

Funding acquisition - SFW, NPG, KZ, XJW.

## Supplementary Figures

**Figure S1.**
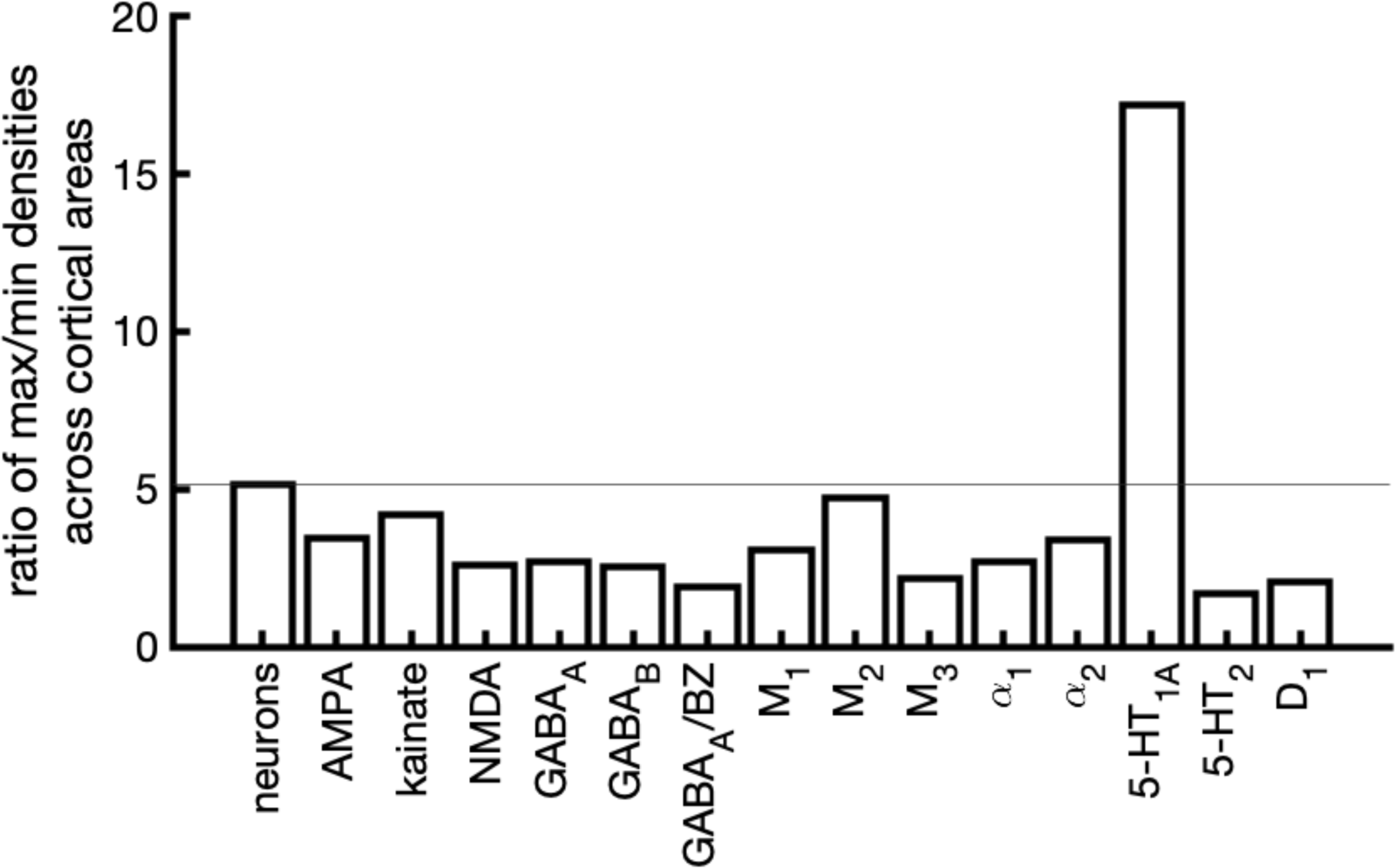
A steep gradient of 5-HT_1A_ receptors in macaque cortex. The ratio of densities for neurons and each receptor type, formed by the area with maximum density divided by the area with minimal density.

**Figure S2.**
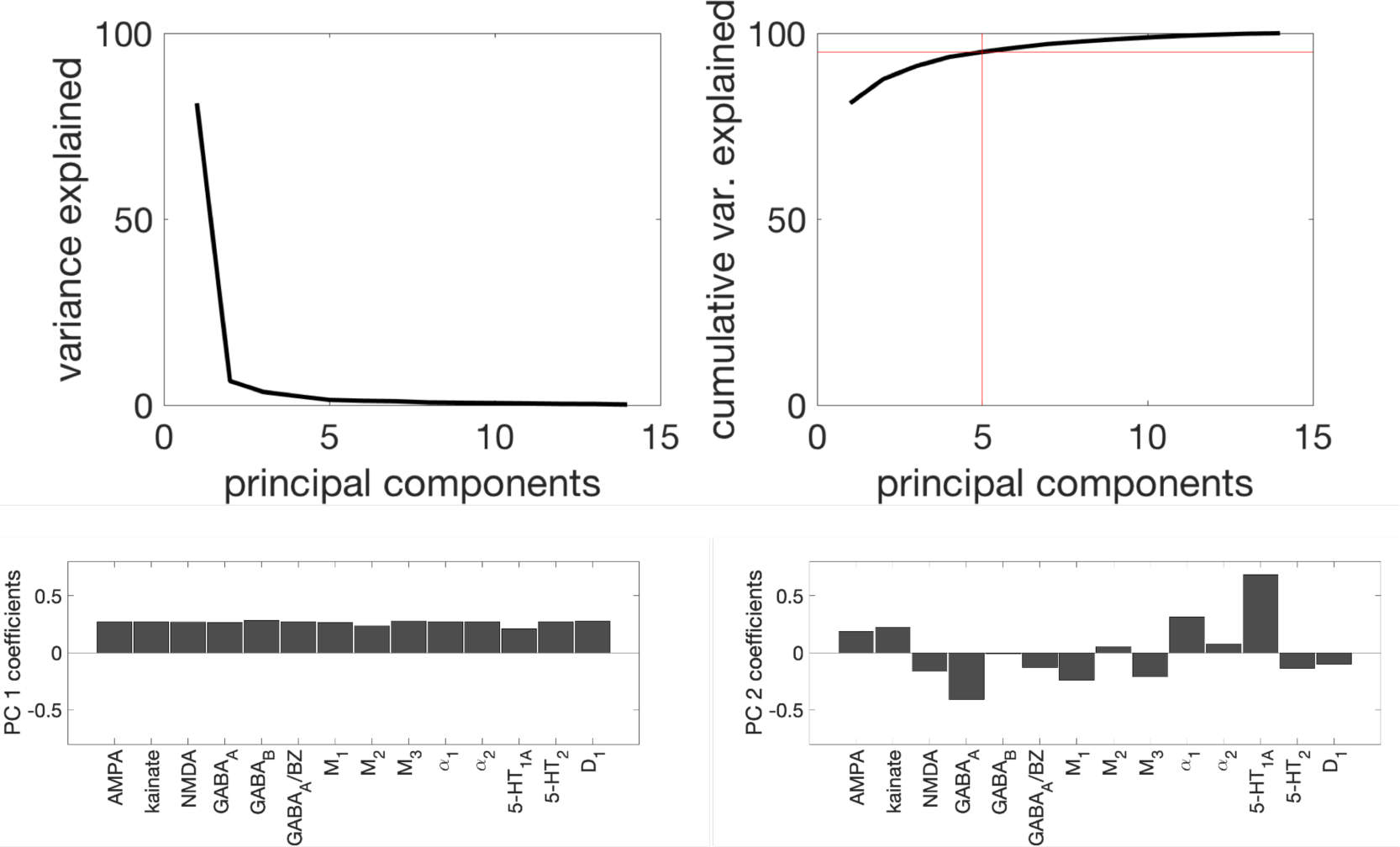
Principal components of the receptor data. Top. The variance explained (left) and cumulative variance explained (right) of each principal component. The top five principal components explained 95% of the variance in the data. The principal component coefficients for each receptor type contributing to principal component 1 (left) and principal component 2 (right).

**Figure S3.**
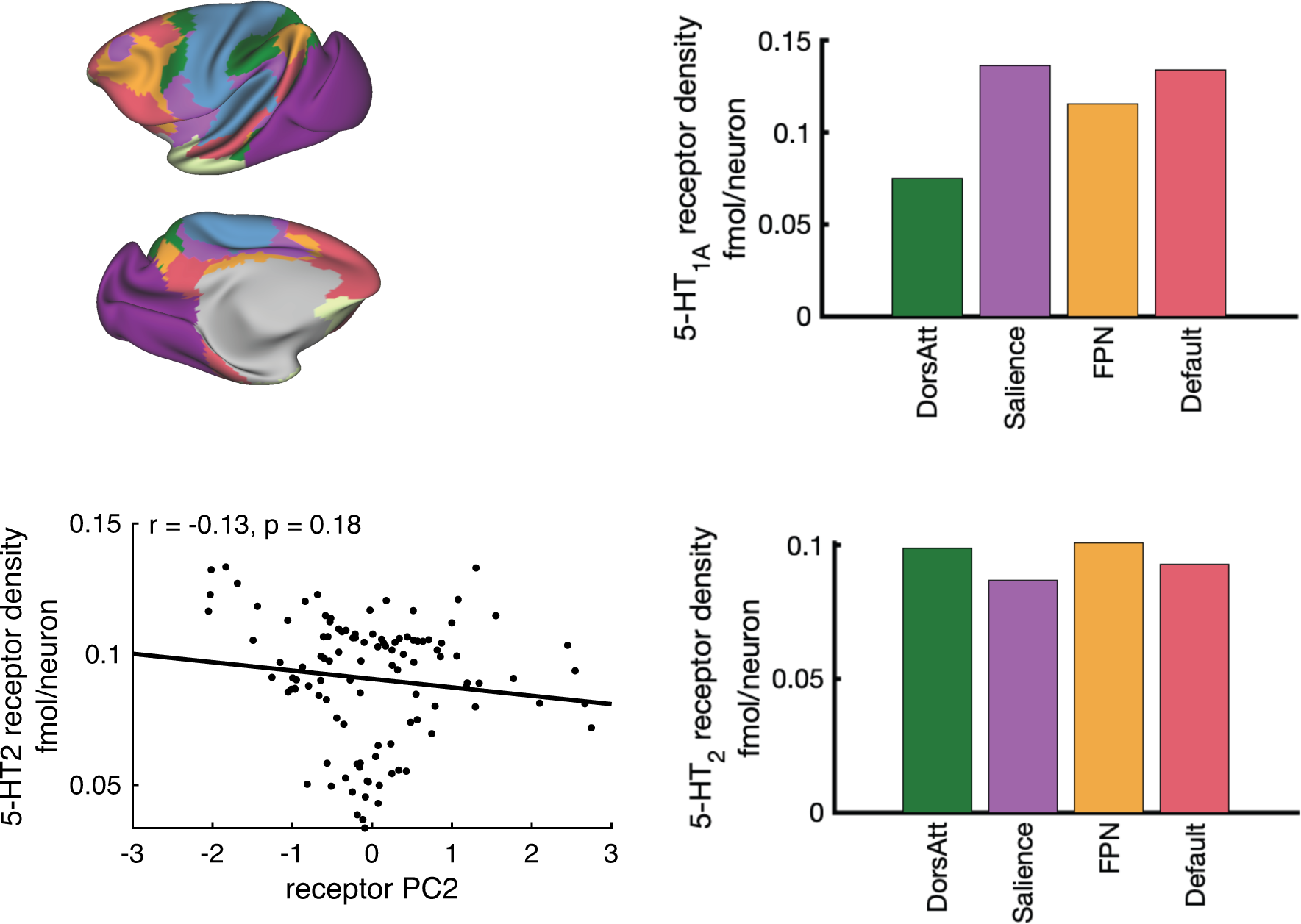
The serotonin 5-HT_1A_ and 5-HT_2_ receptor density in higher cognitive networks. Top-left. The Yeo-Krienen et al cognitive networks in macaque cortex). Right. The mean receptor density for 5-HT_1A_ receptors and 5-HT_2_ receptors in the higher cognitive networks. Note, both receptor types were weakly expressed (per neuron) in the sensory networks. Bottom-left. There was no significant correlation between 5-HT2 expression and receptor PC2.

**Supplementary Table 1.** Overlap of regions of the Julich Macaque Brain Atlas with the 7 cognitive networks of Yeo, Krienen et al in the macaque cortex. Overlap is expressed as a fraction.

|  | Visual | SomMot | DorsAtt | Salience | Limbic | FPN | Default |
| --- | --- | --- | --- | --- | --- | --- | --- |
| 10d | 0 | 0 | 0 | 0 | 0.34 | 0 | 0.66 |
| 10md | 0 | 0 | 0 | 0 | 0 | 0 | 1 |
| 10mv | 0 | 0 | 0 | 0 | 0.11 | 0 | 0.89 |
| 10o | 0 | 0 | 0 | 0 | 0.28 | 0 | 0.72 |
| 11l | 0 | 0 | 0 | 0 | 0.83 | 0.17 | 0 |
| 11m | 0 | 0 | 0 | 0 | 0.9 | 0 | 0.1 |
| 12l | 0 | 0 | 0 | 0 | 0 | 0 | 1 |
| 12m | 0 | 0 | 0 | 0.05 | 0 | 0.7 | 0.25 |
| 12o | 0 | 0 | 0 | 0.02 | 0.24 | 0.03 | 0.7 |
| 12r | 0 | 0 | 0 | 0 | 0.26 | 0.14 | 0.6 |
| 13b | 0 | 0 | 0 | 0 | 0.99 | 0 | 0.01 |
| 13l | 0 | 0 | 0 | 0 | 1 | 0 | 0 |
| 13m | 0 | 0 | 0 | 0 | 1 | 0 | 0 |
| 14r | 0 | 0 | 0 | 0 | 0.78 | 0 | 0.22 |
| 44 | 0 | 0 | 0.45 | 0 | 0 | 0.34 | 0.21 |
| 45A | 0 | 0 | 0 | 0 | 0 | 0.25 | 0.75 |
| 45B | 0 | 0 | 0 | 0 | 0 | 0.62 | 0.38 |
| 4a | 0 | 0.87 | 0.1 | 0.03 | 0 | 0 | 0 |
| 4m | 0 | 1 | 0 | 0 | 0 | 0 | 0 |
| 4p | 0 | 0.99 | 0 | 0.01 | 0 | 0 | 0 |
| 8Ad | 0 | 0 | 0 | 0.27 | 0 | 0.72 | 0.01 |
| 8Av | 0 | 0 | 0.01 | 0 | 0 | 0.98 | 0.01 |
| 8Bd | 0 | 0 | 0 | 0 | 0 | 0 | 1 |
| 8Bm | 0 | 0 | 0 | 0 | 0 | 0 | 1 |
| 8Bs | 0 | 0 | 0 | 0.25 | 0 | 0.35 | 0.4 |
| 9d | 0 | 0 | 0 | 0.14 | 0 | 0.08 | 0.78 |
| 9m | 0 | 0 | 0 | 0 | 0 | 0 | 1 |
| a46d | 0 | 0 | 0 | 0 | 0 | 0.53 | 0.47 |
| a46dr | 0 | 0 | 0 | 0 | 0.05 | 0.27 | 0.67 |
| a46v | 0 | 0 | 0 | 0 | 0 | 0.79 | 0.21 |
| a46vr | 0 | 0 | 0 | 0 | 0.07 | 0.25 | 0.68 |
| F2d | 0 | 0.21 | 0.48 | 0.07 | 0 | 0.24 | 0 |
| F2v | 0 | 0.02 | 0.5 | 0 | 0 | 0.18 | 0.3 |
| F3 | 0 | 0.3 | 0 | 0.42 | 0 | 0.06 | 0.22 |
| F4d | 0 | 0.36 | 0.63 | 0.02 | 0 | 0 | 0 |
| F4s | 0 | 0 | 0.68 | 0 | 0 | 0.32 | 0 |
| F4v | 0 | 0.1 | 0 | 0.9 | 0 | 0 | 0 |
| F5d | 0 | 0 | 0 | 0.67 | 0 | 0.22 | 0.12 |
| F5s | 0 | 0 | 0.44 | 0.12 | 0 | 0.13 | 0.31 |
| F5v | 0 | 0 | 0 | 0.35 | 0 | 0.08 | 0.57 |
| F6 | 0 | 0 | 0 | 0 | 0 | 0.23 | 0.77 |
| F7d | 0 | 0 | 0 | 0 | 0 | 0.18 | 0.82 |
| F7i | 0 | 0 | 0 | 0 | 0 | 0.11 | 0.89 |
| F7s | 0 | 0 | 0 | 0 | 0 | 0.08 | 0.92 |
| p46d | 0 | 0 | 0 | 0 | 0 | 1 | 0 |
| p46dr | 0 | 0 | 0 | 0.58 | 0 | 0.42 | 0 |
| p46v | 0 | 0 | 0 | 0 | 0 | 1 | 0 |
| p46vr | 0 | 0 | 0 | 0 | 0 | 0.67 | 0.33 |
| 1 | 0 | 0.92 | 0 | 0.08 | 0 | 0 | 0 |
| 2 | 0 | 0.98 | 0 | 0.02 | 0 | 0 | 0 |
| 3al | 0 | 0.81 | 0 | 0.17 | 0 | 0.01 | 0.02 |
| 3am | 0 | 1 | 0 | 0 | 0 | 0 | 0 |
| 3bl | 0 | 0.93 | 0 | 0.06 | 0 | 0 | 0 |
| 3bm | 0 | 1 | 0 | 0 | 0 | 0 | 0 |
| AIP | 0 | 0.05 | 0.95 | 0 | 0 | 0 | 0 |
| DP | 0.42 | 0 | 0.55 | 0 | 0 | 0.03 | 0 |
| LIPd | 0 | 0 | 0.82 | 0 | 0 | 0.18 | 0 |
| LIPv | 0 | 0 | 1 | 0 | 0 | 0 | 0 |
| LOP | 0 | 0 | 1 | 0 | 0 | 0 | 0 |
| MIPd | 0 | 0.47 | 0.53 | 0 | 0 | 0 | 0 |
| MIPv | 0 | 0.06 | 0.94 | 0 | 0 | 0 | 0 |
| Opt | 0 | 0 | 0.29 | 0 | 0 | 0.71 | 0 |
| PEc | 0 | 0.31 | 0.69 | 0 | 0 | 0 | 0 |
| PEci | 0 | 1 | 0 | 0 | 0 | 0 | 0 |
| PEipe | 0 | 0.79 | 0.21 | 0 | 0 | 0 | 0 |
| PEipi | 0 | 0.08 | 0.92 | 0 | 0 | 0 | 0 |
| PEl | 0 | 0.92 | 0.08 | 0 | 0 | 0 | 0 |
| PEla | 0 | 1 | 0 | 0 | 0 | 0 | 0 |
| PF | 0 | 0.23 | 0.43 | 0.34 | 0 | 0 | 0 |
| PFG | 0 | 0 | 1 | 0 | 0 | 0 | 0 |
| PFop | 0 | 0.36 | 0.06 | 0.58 | 0 | 0 | 0 |
| PG | 0 | 0 | 0.75 | 0 | 0 | 0.25 | 0 |
| PGm | 0 | 0 | 0.3 | 0.5 | 0 | 0.02 | 0.18 |
| PGop | 0 | 0 | 0.65 | 0.31 | 0 | 0.04 | 0 |
| PIP | 0.02 | 0 | 0.98 | 0 | 0 | 0 | 0 |
| PPt | 0 | 0 | 0.02 | 0 | 0 | 0.98 | 0 |
| TSA | 0 | 1 | 0 | 0 | 0 | 0 | 0 |
| V6Adl | 0 | 0 | 1 | 0 | 0 | 0 | 0 |
| V6Adm | 0 | 0 | 0.31 | 0 | 0 | 0.4 | 0.29 |
| V6Avl | 0 | 0 | 0.95 | 0 | 0 | 0.05 | 0 |
| V6Avm | 0.01 | 0 | 0.01 | 0 | 0 | 0.53 | 0.45 |
| V6l | 0.57 | 0 | 0.43 | 0 | 0 | 0 | 0 |
| V6m | 0.72 | 0 | 0.04 | 0 | 0 | 0.23 | 0 |
| VIP | 0 | 0.02 | 0.98 | 0 | 0 | 0 | 0 |
| V1 | 1 | 0 | 0 | 0 | 0 | 0 | 0 |
| V2d | 0.92 | 0 | 0 | 0 | 0 | 0 | 0.08 |
| V2v | 0.99 | 0 | 0 | 0 | 0 | 0 | 0 |
| V3A | 0.95 | 0 | 0.05 | 0 | 0 | 0 | 0 |
| V3d | 0.92 | 0 | 0.08 | 0 | 0 | 0 | 0 |
| V3v | 1 | 0 | 0 | 0 | 0 | 0 | 0 |
| V4dl | 0.96 | 0 | 0.04 | 0 | 0 | 0 | 0 |
| V4dm | 0.24 | 0 | 0.76 | 0 | 0 | 0 | 0 |
| V4v | 0.92 | 0 | 0 | 0 | 0 | 0 | 0.06 |
| 23c | 0 | 0.96 | 0 | 0.04 | 0 | 0 | 0 |
| 23d | 0 | 0.33 | 0 | 0.55 | 0 | 0.06 | 0.06 |
| 24ab | 0 | 0 | 0 | 0 | 0 | 0 | 1 |
| 24c | 0 | 0 | 0 | 0 | 0 | 0 | 1 |
| 24d | 0 | 0.26 | 0 | 0.68 | 0 | 0.06 | 0 |
| 25 | 0 | 0 | 0 | 0 | 0.74 | 0 | 0.26 |
| 29/30 | 0 | 0 | 0 | 0.02 | 0 | 0 | 0.5 |
| 31 | 0 | 0.63 | 0.04 | 0.33 | 0 | 0 | 0 |
| a24ab | 0 | 0 | 0 | 0.29 | 0 | 0.17 | 0.54 |
| a24c | 0 | 0 | 0 | 0.06 | 0 | 0.38 | 0.56 |
| d23ab | 0 | 0 | 0 | 0.51 | 0 | 0.02 | 0.48 |
| p24ab | 0 | 0.01 | 0 | 0.94 | 0 | 0.06 | 0 |
| p32 | 0 | 0 | 0 | 0 | 0 | 0 | 1 |
| s32 | 0 | 0 | 0 | 0 | 0.08 | 0 | 0.92 |
| v23ab | 0 | 0 | 0 | 0 | 0 | 0 | 0.96 |
| MT | 0.07 | 0.09 | 0.21 | 0.11 | 0 | 0.09 | 0.44 |

**Supplementary Table 2:**
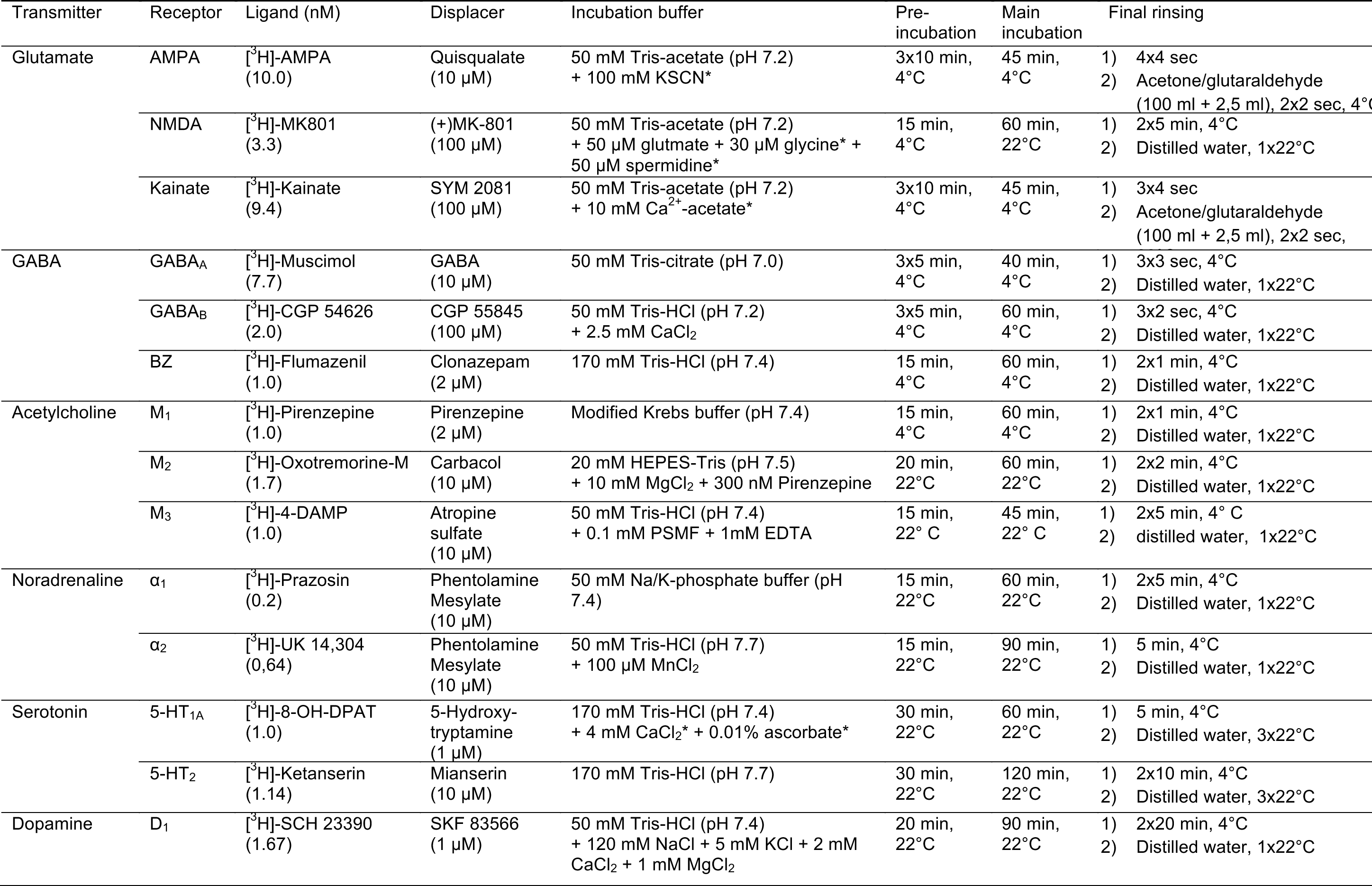
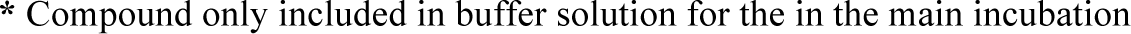
Incubation protocols.

| Transmitter | Receptor | Ligand (nM) | Displacer | Incubation buffer | Pre-incubation | Main incubation | Final rinsing |
| --- | --- | --- | --- | --- | --- | --- | --- |
| Glutamate | AMPA | [<sup>3</sup>H]-AMPA (10.0) | Quisqualate (10 µM) | 50 mM Tris-acetate (pH 7.2) + 100 mM KSCN* | 3x10 min, 4°C | 45 min, 4°C | 1) 4x4 sec<br>2) Acetone/glutaraldehyde (100 ml + 2,5 ml), 2x2 sec, 4°C |
|  | NMDA | [<sup>3</sup>H]-MK801 (3.3) | (+)MK-801 (100 µM) | 50 mM Tris-acetate (pH 7.2) + 50 µM glutamate + 30 µM glycine* + 50 µM spermidine* | 15 min, 4°C | 60 min, 22°C | 1) 2x5 min, 4°C<br>2) Distilled water, 1x22°C |
|  | Kainate | [<sup>3</sup>H]-Kainate (9.4) | SYM 2081 (100 µM) | 50 mM Tris-acetate (pH 7.2) + 10 mM Ca<sup>2+</sup>-acetate* | 3x10 min, 4°C | 45 min, 4°C | 1) 3x4 sec<br>2) Acetone/glutaraldehyde (100 ml + 2,5 ml), 2x2 sec, |
| GABA | GABA<sub>A</sub> | [<sup>3</sup>H]-Muscimol (7.7) | GABA (10 µM) | 50 mM Tris-citrate (pH 7.0) | 3x5 min, 4°C | 40 min, 4°C | 1) 3x3 sec, 4°C<br>2) Distilled water, 1x22°C |
|  | GABA<sub>B</sub> | [<sup>3</sup>H]-CGP 54626 (2.0) | CGP 55845 (100 µM) | 50 mM Tris-HCl (pH 7.2) + 2.5 mM CaCl<sub>2</sub> | 3x5 min, 4°C | 60 min, 4°C | 1) 3x2 sec, 4°C<br>2) Distilled water, 1x22°C |
|  | BZ | [<sup>3</sup>H]-Flumazenil (1.0) | Clonazepam (2 µM) | 170 mM Tris-HCl (pH 7.4) | 15 min, 4°C | 60 min, 4°C | 1) 2x1 min, 4°C<br>2) Distilled water, 1x22°C |
| Acetylcholine | M<sub>1</sub> | [<sup>3</sup>H]-Pirenzepine (1.0) | Pirenzepine (2 µM) | Modified Krebs buffer (pH 7.4) | 15 min, 4°C | 60 min, 4°C | 1) 2x1 min, 4°C<br>2) Distilled water, 1x22°C |
|  | M<sub>2</sub> | [<sup>3</sup>H]-Oxotremorine-M (1.7) | Carbacol (10 µM) | 20 mM HEPES-Tris (pH 7.5) + 10 mM MgCl<sub>2</sub> + 300 nM Pirenzepine | 20 min, 22°C | 60 min, 22°C | 1) 2x2 min, 4°C<br>2) Distilled water, 1x22°C |
|  | M<sub>3</sub> | [<sup>3</sup>H]-4-DAMP (1.0) | Atropine sulfate (10 µM) | 50 mM Tris-HCl (pH 7.4) + 0.1 mM PSMF + 1mM EDTA | 15 min, 22° C | 45 min, 22° C | 1) 2x5 min, 4° C<br>2) distilled water, 1x22°C |
| Noradrenaline | α<sub>1</sub> | [<sup>3</sup>H]-Prazosin (0.2) | Phentolamine Mesylate (10 µM) | 50 mM Na/K-phosphate buffer (pH 7.4) | 15 min, 22°C | 60 min, 22°C | 1) 2x5 min, 4°C<br>2) Distilled water, 1x22°C |
|  | α<sub>2</sub> | [<sup>3</sup>H]-UK 14,304 (0,64) | Phentolamine Mesylate (10 µM) | 50 mM Tris-HCl (pH 7.7) + 100 µM MnCl<sub>2</sub> | 15 min, 22°C | 90 min, 22°C | 1) 5 min, 4°C<br>2) Distilled water, 1x22°C |
| Serotonin | 5-HT<sub>1A</sub> | [<sup>3</sup>H]-8-OH-DPAT (1.0) | 5-Hydroxy-tryptamine (1 µM) | 170 mM Tris-HCl (pH 7.4) + 4 mM CaCl<sub>2</sub>* + 0.01% ascorbate* | 30 min, 22°C | 60 min, 22°C | 1) 5 min, 4°C<br>2) Distilled water, 3x22°C |
|  | 5-HT<sub>2</sub> | [<sup>3</sup>H]-Ketanserin (1.14) | Mianserin (10 µM) | 170 mM Tris-HCl (pH 7.7) | 30 min, 22°C | 120 min, 22°C | 1) 2x10 min, 4°C<br>2) Distilled water, 3x22°C |
| Dopamine | D<sub>1</sub> | [<sup>3</sup>H]-SCH 23390 (1.67) | SKF 83566 (1 µM) | 50 mM Tris-HCl (pH 7.4) + 120 mM NaCl + 5 mM KCl + 2 mM CaCl<sub>2</sub> + 1 mM MgCl<sub>2</sub> | 20 min, 22°C | 90 min, 22°C | 1) 2x20 min, 4°C<br>2) Distilled water, 1x22°C |

\* Compound only included in buffer solution for the in the main incubation

